# Climate-induced forest dieback drives compositional change in insect communities that is concentrated amongst rare species

**DOI:** 10.1101/2021.04.21.440751

**Authors:** Lucas Sire, Paul Schmidt Yáñez, Cai Wang, Annie Bézier, Béatrice Courtial, Jérémy Cours, Diego Fontaneto, Laurent Larrieu, Christophe Bouget, Simon Thorn, Jörg Müller, Douglas W. Yu, Michael T. Monaghan, Elisabeth A. Herniou, Carlos Lopez-Vaamonde

**Author notes:** corresponding author (L. Sire).

## Abstract

Marked decline in insect species richness, abundance and biomass have recently been quantified in Europe. We metabarcoded 224 Malaise-trap samples to investigate whether drought-induced forest dieback and subsequent salvage logging have an impact on flying insects (*ca*. 3000 insect species) in silver fir Pyrenean forests. We found no evidence that climate-induced forest dieback impacted species richness of flying insects but revealed compositional turnover patterns consistent with those seen during natural forest succession, given that the key covariates explaining compositional variation were canopy openness versus microhabitat diversity and deadwood amount at local and landscape scales, respectively. Importantly, most change was driven by rare species. In contrast, observed levels of salvage logging did not explain change in species richness or composition. Hence, although forest dieback appears to cause changes in species assemblages mimicking natural forest succession, it also increases the risk of catastrophic loss of rare species through homogenization of environmental conditions.

## Introduction

Insects are vital components of biodiversity, providing important ecosystem services such as pollination and pests regulation, while also performing disservices as disease vectors and plant pests^1^. Global changes, including those of climate, land use and land cover, can lead to degradation and habitat loss, chemical and light pollution or invasive species. These changes have caused major drops in biomass, abundance and species richness of insects^2^, and this accelerating decline has become a major cause for concern. Yet, understanding the relative impacts of these drivers underlying insect decline and its propensity is complex^3^.

Forests are thought to act as buffer zones from rapid anthropogenic changes in adjacent areas, and contribute to biodiversity conservation by providing refuge for rich insect communities^4^. However, forests are increasingly suffering from climate-induced tree dieback, regardless of protection measures^5^, which could have long term consequences for forest biodiversity. Tree diebacks largely result from more frequent, intense and longer droughts, especially in Europe^6^. These can induce tree mortality especially if they promote insect pests and pathogen outbreaks. Some insect species benefit from warmer temperatures (*i.e.* mild winters) allowing distribution range expansion and population outbreaks, which can cause massive forest diebacks^7^. Bioclimatic models predict that the extent, severity and duration of droughts will increase as a result of climate change, notably in temperate and Mediterranean areas^8^.

Tree diebacks can cause major structural changes such as an increase in canopy openness, reduced foliage density, which in turn increases light availability, potentially changing the community structure of understory plants and their associated herbivorous insects^9^. While insect decline is independent from forest protection status, it has been presented as lower in terms of species richness for plots undergoing dieback^2^. Indeed, tree dieback can also increase the availability of nutrients stored in vital trees (*i.e*. deadwood), sunlight and microhabitats, increasing species richness for multiple insect groups including bees, wasps, hoverflies, saproxylic beetles, as well as multiple red-listed insect taxa following insect pest outbreaks^10^, or canopy-dwelling beetles in declining oak forests^9^. However, no significant change in species richness of ground-dwelling carabids and spiders has been associated with climate-induced dieback in a beech-dominated forest^11^. Meanwhile, forest management practices, like salvage logging (*i.e.* cutting down dying trees to salvage their timber value), can also have contrasting effects on biodiversity^12^. For instance, species richness of saproxylic insects increased with bark beetle outbreaks^10^, but decreased with deadwood harvesting^13^. Even though specific to each of these well-studied taxa previously stated, species richness in general appears to be boosted by tree diebacks, while salvage logging seems to have no effect overall^14^.

The identification of samples down to species level is important in ecological studies because adaptive response to disturbances can be highly species-specific^15^. Moreover, these studies on insect biodiversity response to forest disturbances are mainly based on species richness, which has been used as a surrogate for ecosystem functionality^16^. However, richness alone is often a poor indicator of biodiversity change compared to ecological guilds changes to the community^17^. Quantifying these changes requires high taxonomic resolution that is often not available. The morphological identification that richness measures are typically derived from are often limited to well-known insect bioindicators that are overrepresented in the literature, hence biasing species richness *per se* and impeding further holistic community-based and ecological studies. Unfortunately, accurate species identification of hyper-diverse insect groups such as dipterans and hymenopterans is difficult because of the taxonomic impediment^18^, often hampering—or limiting to few remarkable groups only—the study of these species-rich taxa despite their ecological importance^19^. The use of DNA barcoding as a tool for species identification^20^ now allows to bypass this taxonomic impediment^21^. As exemplified by Wang *et al.*^22^ targeting poorly described Chinese entomofauna, the use of DNA metabarcoding allows to tackle more comprehensive insect biodiversity studies and facilitates the documenting of all trophic guilds and their responses to forest disturbances. The authors suggested that forest diebacks induced by bark beetle outbreaks could drive a transition from homogenous plantations to more biodiversity-friendly heterogenous forests^22^.

The response to forest dieback and subsequent salvage logging varies among taxa and functional groups. Floricolous species are expected to increase in diversity and abundance from tree dieback due to greater canopy openness^15^. Similarly, parasitoids may benefit from temperature rises and associated tree diebacks as the more complex a forest is the more diverse the parasitoid communities are^23^, but may also suffer from poorer host quality, especially of sap-feeders directly affected by tree health under drought conditions^24^. Forests undergoing diebacks are thus expected to induce guild-specific responses, which may have contrasting effects on species richness of each insect groups depending on their respective ecology. A comprehensive approach on sampling and identification of taxa is therefore needed to improve our understanding of the effects of forest disturbances on insect biodiversity.

Here, we study insect diversity in montane Pyrenean forests dominated by silver fir (*Abies alba* Mill), a conifer species sensitive to drought^25^ that suffered severe climate-induced diebacks after a heatwave in 2003^26^. Currently, forest management practices often implement salvage logging at the first signs of dieback^26^. Despite the high conservation value of Pyrenean silver fir forests, the effects of dieback and salvage logging on associated insect fauna remains unknown. To address this gap, we studied the communities of aerial insects over 56 silver fir forest plots (Supplementary Fig. 1) that varied in dieback level and management practices. From late spring to early autumn 2017, insects were mass-trapped monthly using Malaise traps, resulting in 224 samples that we analyzed using DNA metabarcoding of a fragment of the mitochondrial COI gene region. Based on BOLD DNA reference libraries^27^, taxonomy of the recovered Molecular Operational Taxonomic Units (MOTUs) was assessed and broad ecological functions (*i.e.* floricolous and parasitoid insects) were attributed using family-level information from published literature whenever possible. We used generalized linear models to assess whether dieback levels and salvage logging influence the structure and diversity of forest insect communities and functional guilds. We hypothesized that levels of forest dieback and salvage logging change species composition of local insect communities, which is seen as turnover in community structure across sites with different disturbances. We also hypothesized that functional guilds respond differently to forest dieback intensity; in particular the diversity of parasitoids and floricolous insects in areas of high forest dieback. As we found no change in species richness but variations in community compositions across dieback levels and management practices, we performed zeta order analyses^28, 29^ to further investigate the dataset structure and identify the nature of these community changes, as well as the species compositional and functional turnover. We also used zeta analyses in multi-site generalized dissimilarity modelling (zeta.msgdm)^30^ to study the impact of environmental features (*i.e.* geographic distance to the nearest plot, altitude, canopy openness, total amount of deadwood, basal area, tree diversity, density of very large trees and Tree-related Microhabitats (TreM) diversity) in driving these community structural changes and further discuss their associated consequences for forest management and conservation. Finally, we used the fourth-corner model^31^ to highlight winning and losing insects to forest disturbances as well as idiosyncratic responses of insect orders and functional groups.

## Results

### Diverse insect community with high temporal and spatial turnover

Metabarcoding recovered thousands of locally rare species and unnamed “dark taxa” (*i.e.* specie-level ‘invertebrate’ taxa in DNA reference libraries without a species name assigned and/or species-rich taxa, often small in body size and under-described)^32^. From the 224 Malaise trap samples, we recovered 2972 MOTUs, with iNEXT extrapolation estimating a total richness of ∼4000 MOTUs and species accumulation curves reaching a plateau (Fig. 1A, Supplementary Fig. 2). Our 4-month sampling effort (from late spring to early autumn) thus captured *ca.* 75% of the total Malaise-trappable insect diversity in the sampled areas allowing us to reach ∼90% sample coverage (Fig. 1A–C). Applying a threshold of 97% sequence similarity for species-level discrimination and keeping unambiguous taxonomic matches for higher taxonomic ranks, 100%, 84.8%, 75.4% and 52.4% (1558) of the total 2972 insect MOTUs could be assigned to order, family, genus and species names, respectively (Supplementary Table I). Two orders, Diptera and Hymenoptera, together represented 73% of all MOTUs, as expected with Malaise-trapped samples (Fig. 2A), with 641 (49.7%) and 371 MOTUs (41.4%) identified to species, respectively (Supplementary Fig. 4). As with the whole dataset (Fig. 1A), the accumulation curves of the five most represented insect orders approached asymptotes (Fig. 2B).

**Fig. 1:**
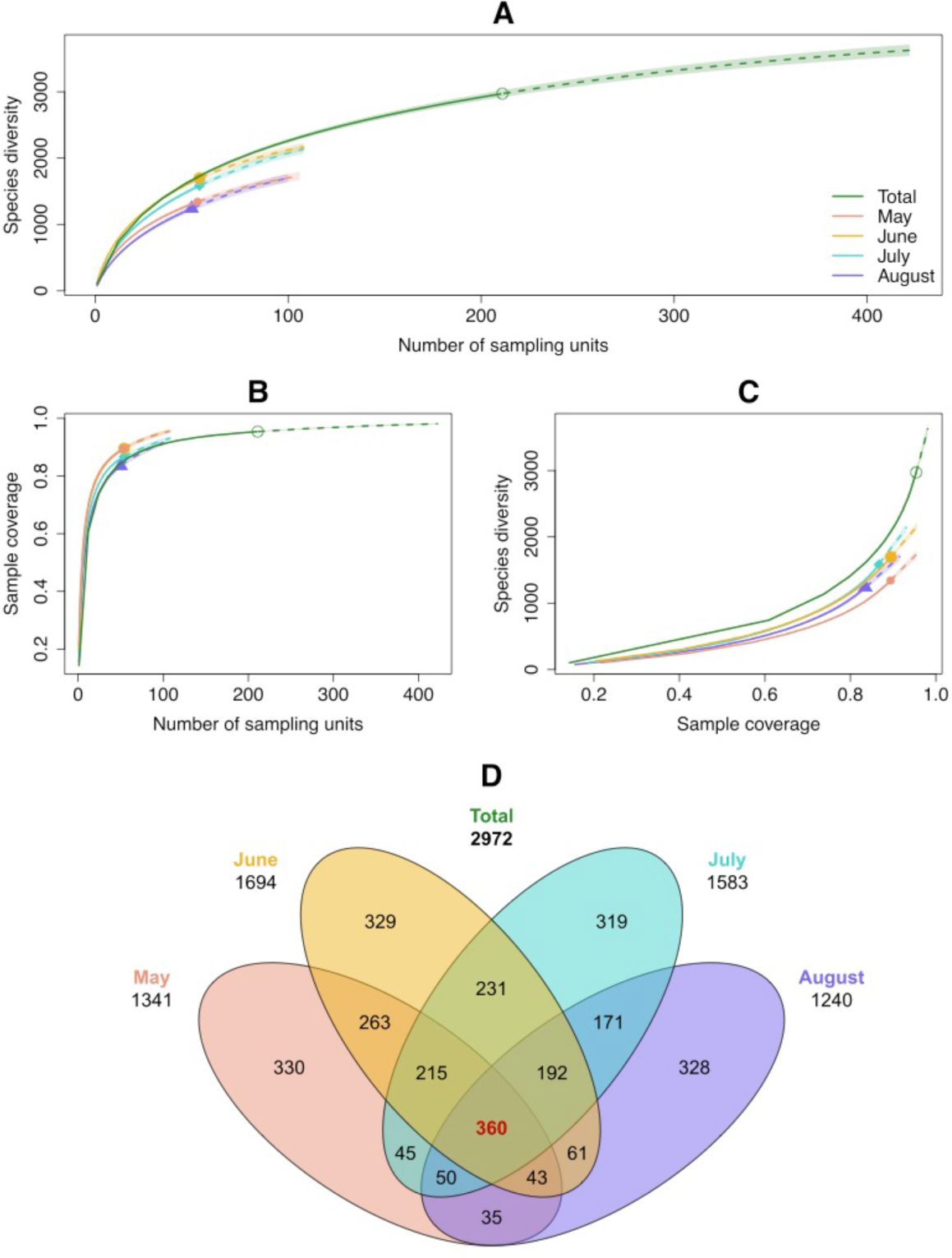
iNext species accumulation curves for Malaise trap samples and temporal turnover. **(A)** Species richness (MOTUs) per sampling unit recovered through metabarcoding of Malaise trap samples (solid lines). Incidence frequency-based species richness extrapolations (dashed lines) were performed over the total sampling period (from May 15 to September 15, 2017) (green curve) as well as for the following four-weeks temporal sampling categories: May, June, July, August to assess the potential variations in both sampling effort and species richness (pink, yellow, cyan and violet curves, respectively). Sampling units are the total number of Malaise trap samples available per temporal category. Shaded area represents 95% confidence interval. **(B)** Sample coverage (sampling efficiency of sampling units to recover the expected biodiversity) per number of sampling units; (**C**) Species diversity (MOTUs) per sample coverage. Accumulation curves indicated that the 224 sampling units used in the present study allowed a 90% sampling coverage of the given area and sampling period and allowed us to recover nearly 75% of the total Malaise-trappable diversity. Estimations gave around 4000 trappable species and the need of more than 400 sampling units on-site to recover them. (**D**) Venn diagram showing the species (MOTUs) turnover in the French Pyrenees across the four-month sampling period of the study. Total number of MOTUs found every four weeks is given under the name of each starting month. Number of MOTUs shared between months is given within the diagrams. The high MOTU numbers specific to each month and the comparatively low number of shared MOTUs throughout the entire sampling period (highlighted in red) indicated an important temporal turnover of insect species.

**Fig. 2:**
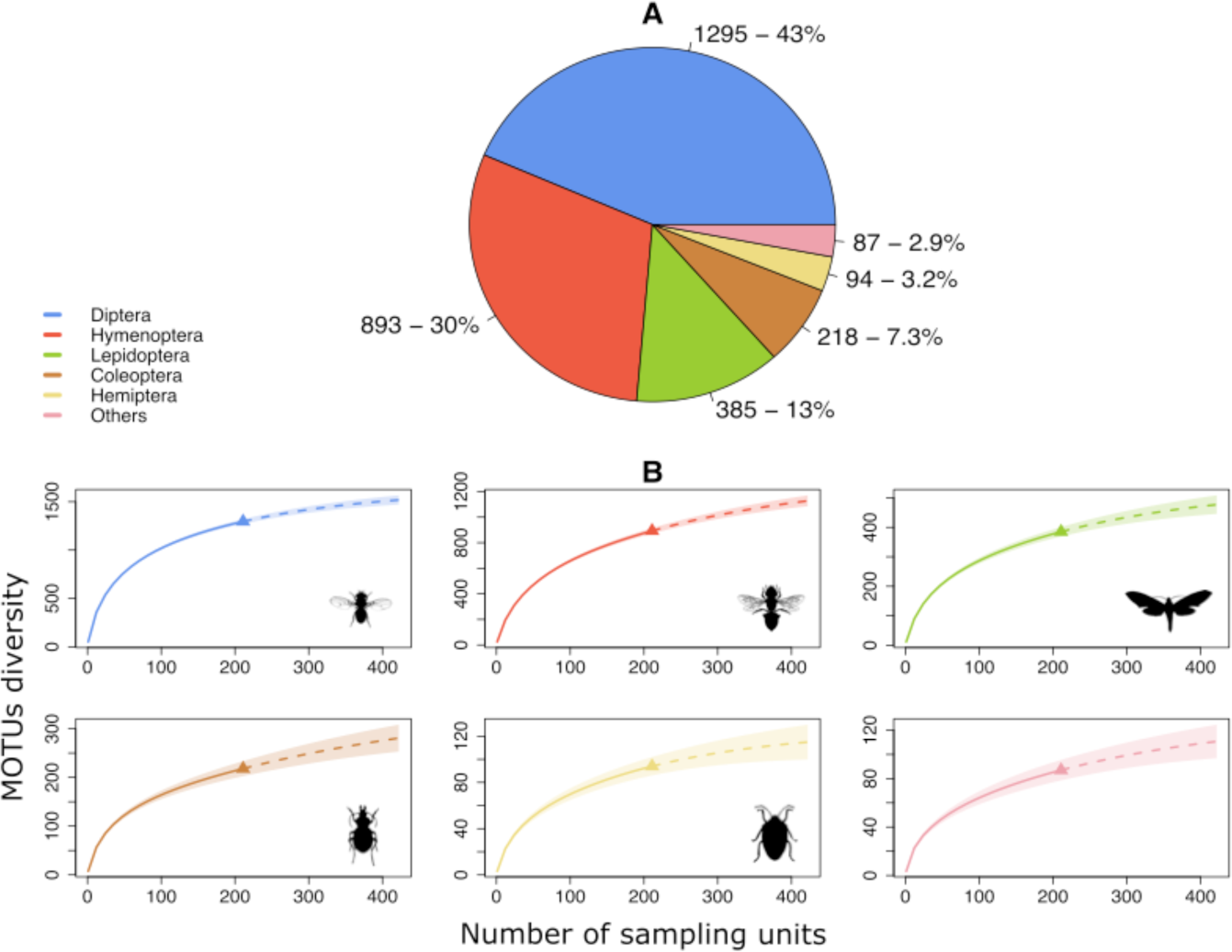
Taxonomic composition and Gamma diversity analysis. (**A**) Pie chart of the 15 insect orders sampled showing the taxonomic affiliation and the respective number of MOTUs recovered, as well as their total proportion. For clarity purposes, only the five orders with highest number of species (Diptera, Hymenoptera, Lepidoptera, Coleoptera and Hemiptera) are represented (in blue, red, green, brown and yellow, respectively), with the 10 remaining orders (Blattodea, Ephemeroptera, Mecoptera, Neuroptera, Orthoptera, Plecoptera, Psocodea, Raphidioptera, Thysanoptera and Trichoptera) clustered within the “Others” category in pink. Representativeness of each insect order in the dataset was consistent with the known bias of Malaise trap sampling, especially toward dipterans and hymenopterans. (**B**) Incidence frequency-based accumulation curves (solid lines) with species richness extrapolations (dashed lines) for each insect order. Shaded area represents 95% confidence interval. As all accumulation curves were nearly plateauing, this indicated that for each represented insect order, almost all Malaise-trappable diversity over the sampling period of had been successfully recovered.

**Table I:** Impact of forest dieback and salvage logging on insect species richness. Generalized linear models for species richness variations of different study groups (*i.e.* total insects, five individual insect orders, and four functional group) compared across respective environmental conditions of diebacks (*i.e.* low, medium and high forest dieback levels) and stand types (*i.e.* healthy, disturbed and salvaged logged). Functional groups (*i.e.* Parasitoids/non-parasitoids, floricolous/non-floricolous insects) were assigned using each MOTU’s taxonomic family. Significance is given by “*” while “*N.S.*” stands for non-significant.

| Studied group | Condition | Degree of freedom | Chi-square | P-value<br>(95% confidence) | Significance |
| --- | --- | --- | --- | --- | --- |
| **Total insects** |  |  |  |  |  |
|  | *Dieback level effect* | 2 | 2.10963 | 0.3483 | N.S. |
|  | *Stand type effect* | 2 | 0.94467 | 0.623 | N.S. |
|  | *District effect* | 1 | 0.51971 | 0.4710 | N.S. |
| **Coleoptera** |  |  |  |  |  |
|  | *Dieback level effect* | 2 | 0.3568 | 0.8365938 | N.S. |
|  | *Stand type effect* | 2 | 0.9060 | 0.6357279 | N.S. |
|  | *District effect* | 1 | 13.9565 | **0.0001871** | **\*\*\*** |
| **Diptera** |  |  |  |  |  |
|  | *Dieback level effect* | 2 | 1.9782 | 0.3719 | N.S. |
|  | *Stand type effect* | 2 | 1.8063 | 0.4053 | N.S. |
|  | *District effect* | 1 | 1.8153 | 0.1779 | N.S. |
| **Hemiptera** |  |  |  |  |  |
|  | *Dieback level effect* | 2 | 0.14194 | 0.9315 | N.S. |
|  | *Stand type effect* | 2 | 1.64021 | 0.4404 | N.S. |
|  | *District effect* | 1 | 1.58658 | 0.2078 | N.S. |
| **Hymenoptera** |  |  |  |  |  |
|  | *Dieback level effect* | 2 | 2.7238 | 0.2562 | N.S. |
|  | *Stand type effect* | 2 | 1.2385 | 0.5384 | N.S. |
|  | *District effect* | 1 | 1.0418 | 0.3074 | N.S. |
| **Lepidoptera** |  |  |  |  |  |
|  | *Dieback level effect* | 2 | 0.74240 | 0.6899 | *N.S.* |
|  | *Stand type effect* | 2 | 0.14966 | 0.9279 | *N.S.* |
|  | *District effect* | 1 | 0.92520 | 0.3361 | *N.S.* |
| **Floricolous** |  |  |  |  |  |
|  | *Dieback level effect* | 2 | 2.33042 | 0.3119 | *N.S.* |
|  | *Stand type effect* | 2 | 0.37240 | 0.8301 | *N.S.* |
|  | *District effect* | 1 | 0.57194 | 0.4495 | *N.S.* |
| **Non-floricolous** |  |  |  |  |  |
|  | *Dieback level effect* | 2 | 2.10039 | 0.3499 | *N.S.* |
|  | *Stand type effect* | 2 | 1.34128 | 0.5114 | *N.S.* |
|  | *District effect* | 1 | 0.10885 | 0.7415 | *N.S.* |
| **Parasitoids** |  |  |  |  |  |
|  | *Dieback level effect* | 2 | 2.4223 | 0.29785 | *N.S.* |
|  | *Stand type effect* | 2 | 0.7983 | 0.67089 | *N.S.* |
|  | *District effect* | 1 | 3.2229 | 0.007261 | *N.S.* |
| **Non-parasitoids** |  |  |  |  |  |
|  | *Dieback level effect* | 2 | 2.01770 | 0.3646 | *N.S.* |
|  | *Stand type effect* | 2 | 1.13627 | 0.5666 | *N.S.* |
|  | *District effect* | 1 | 0.00031 | 0.9860 | *N.S.* |

We observed a high rate of temporal species turnover across months, as evidenced by only 360 MOTUs (12% of the total MOTUs diversity) being detected in all fourth months (Fig. 1D). Thus, even though our sampling was efficient at recovering high biodiversity, sampling throughout the complete growing season is likely to increase the total diversity occurring in the region (Fig. 1D). Geographic turnover was also observed, with only 45% of the total insect species sampled occurring in both of the sampled districts Aure Valley (Central French Pyrenees) and Sault Plateau (Eastern French Pyrenees) (Supplementary Fig. 1), the remaining 31% and 24% being specific to each, respectively. Additionally, these district-specific taxa were the most rarely caught throughout all the different plots of their respective districts (Supplementary Fig. 3A).

### Forest disturbances influences the composition but not the species richness of insect communities

We grouped sites into three dieback categories (low, medium, high) based on the proportion of drought-affected and dying trees for each plot. Insect species richness was compared across the three dieback categories but also across three defined stand types including management practice (healthy, stands expressing dieback but not salvaged—hereafter ‘disturbed’—, stands expressing dieback and being salvage logged—hereafter shortened as salvaged). No response could be detected in terms of changes in species richness from all insect or functional groups tested, neither for the three levels of forest dieback nor the stand types (Table I). However, a significant difference in species richness was found between the two sampled districts for Coleoptera only, with more species of beetles detected in Sault plateau (Table I).

As the dataset was geographically structured (*i.e.* Aure valley and Sault plateau districts), we analysed the dataset using the nearest-neighbour (NN) sample-selection scheme (Fig. 3). The NN model fitted better to a power-law function than to an exponential one (Fig. 3C, D; AIC_*(NN, Exp)*_ = 0.89, AIC_*(NN, PL)*_ = -122.78), which was consistent with community assembly being driven by niche differentiation over stochastic assembly. We also ran a zeta diversity analysis with the non-geographically structured all-combinations (ALL) sample-selection scheme (Supplementary Fig. 5), and the results were similar but showed a much weaker fit to the power-law function (Supplementary Fig. 5C, D; AIC_(ALL, *Exp*_) = 0.63, AIC(ALL, *PL*) = -20.97). The rapid decline in zeta diversity between zeta orders 2 and 10 indicated that compositional turnover was mainly driven by rare species; few species were shared in 10 or more sites (Fig. 3A; Supplementary Fig. 3B). However, re-visualising the decline curve in Fig. 3A as a zeta retention-rate curve (Fig. 3B) showed that the few species that did occur in ≥10 sites (n = 42 species) were highly prevalent but only two species were being shared by 55 out of 56 total sites. Interestingly, this decline curve also demonstrated the drop of species retention rate at 28 sites—equivalent to the number of sampled plots in each district—, accounting for the strong geographic effect that induced very few shared species between Aure valley and Sault plateau. Re-analysis using the ALL sample-selection scheme produced similar results in shared species but smoothed geographic effects (Supplementary Fig. 5).

**Fig. 3:**
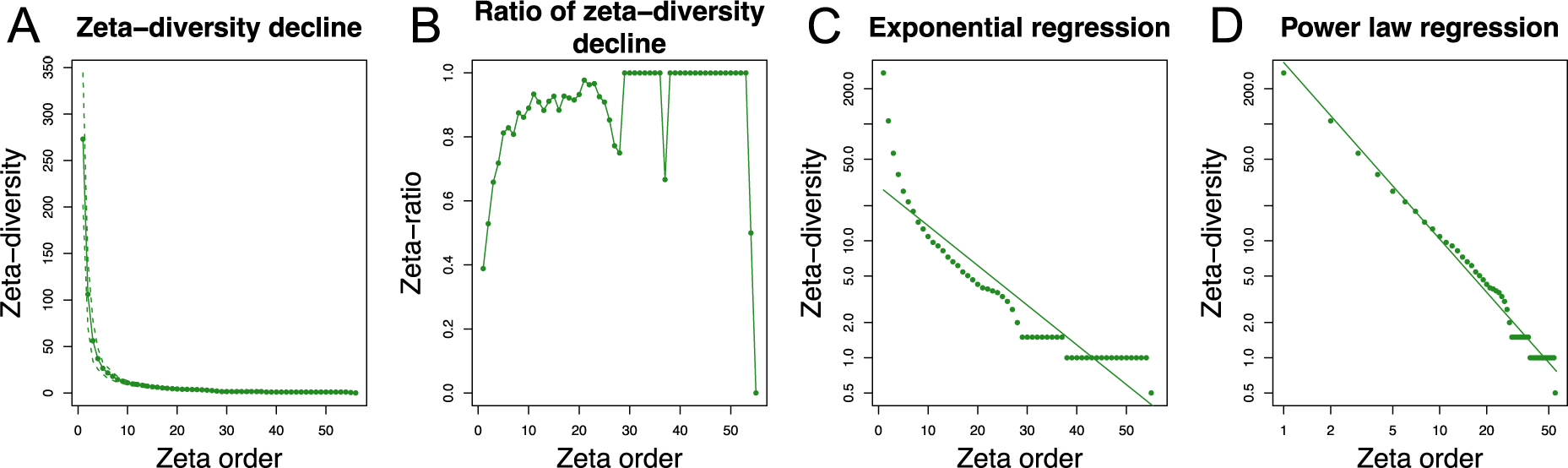
Zeta-diversity decline and model fitting on the insect fauna for nearest-neighbour combinations and assemblages. Zeta-diversity analyses per zeta order (*i.e.* ζi) here referring to plots (from ζ2 to ζ56) for two computing schemes. Representations consider model scheme that computes nearest-neighbour plot combinations and assemblages (NN). (**A, B**) Zeta-diversity decline and 0 to 1 scaled ratio of zeta-diversity decline, representing species shared and species retention rate (*i.e.* the retention probability of common species in the community) across ζi, respectively. Zeta-diversity decline curve (**A**) indicated high heterogeneity in insect diversity across the studied plots as fewer and fewer species were shared between 2–9 plots and almost no common species were being shared when 10 plots or more were considered. Zeta ratio (**B**) highlighted the high prevalence of the few species shared across 2–54 plots which were likely to be sampled, and the drop of species retention rate at zeta order ζ28 corresponded to geographic distance between the two sampled districts. (**C, D**) zeta-diversity model fitting to exponential and power-law regressions, respectively. Fit to exponential regression implies equal and stochastic turnover among species whereas fit to power-law regression as represented here indicates turnover driven by niche differentiation processes and rare species less likely to be found across the whole dataset.

The impacts of drought-induced forest dieback intensities and stand types on community compositional changes were evaluated for the total sampling (Fig. 4A). Overall, forest dieback was found to induce significant changes in insect community assemblages for all tested groups but Coleoptera (Supplementary Table II). However, no significant variation in community composition was found across stand types, hence no effect of salvage logging could be detected (Supplementary Table II). Regardless of dieback intensity and stand type, each insect community assemblage tested significantly differed between districts (Fig. 4B, Supplementary Table II). Each dieback category hosted particular sets of species, yet insect communities of low dieback level plots were more similar to each other and more distinct from those of medium and high dieback level plots (Fig. 4A; Supplementary Fig. 3B). Furthermore, taxa specific to a particular dieback level were mostly rare taxa (*i.e.* taxa with low prevalence) (Supplementary Fig. 3B). Finally, community composition variations found across district were reflected by Sault plateau plots sharing more similar species than those of Aure valley plots (Fig. 4B).

**Fig. 4:**
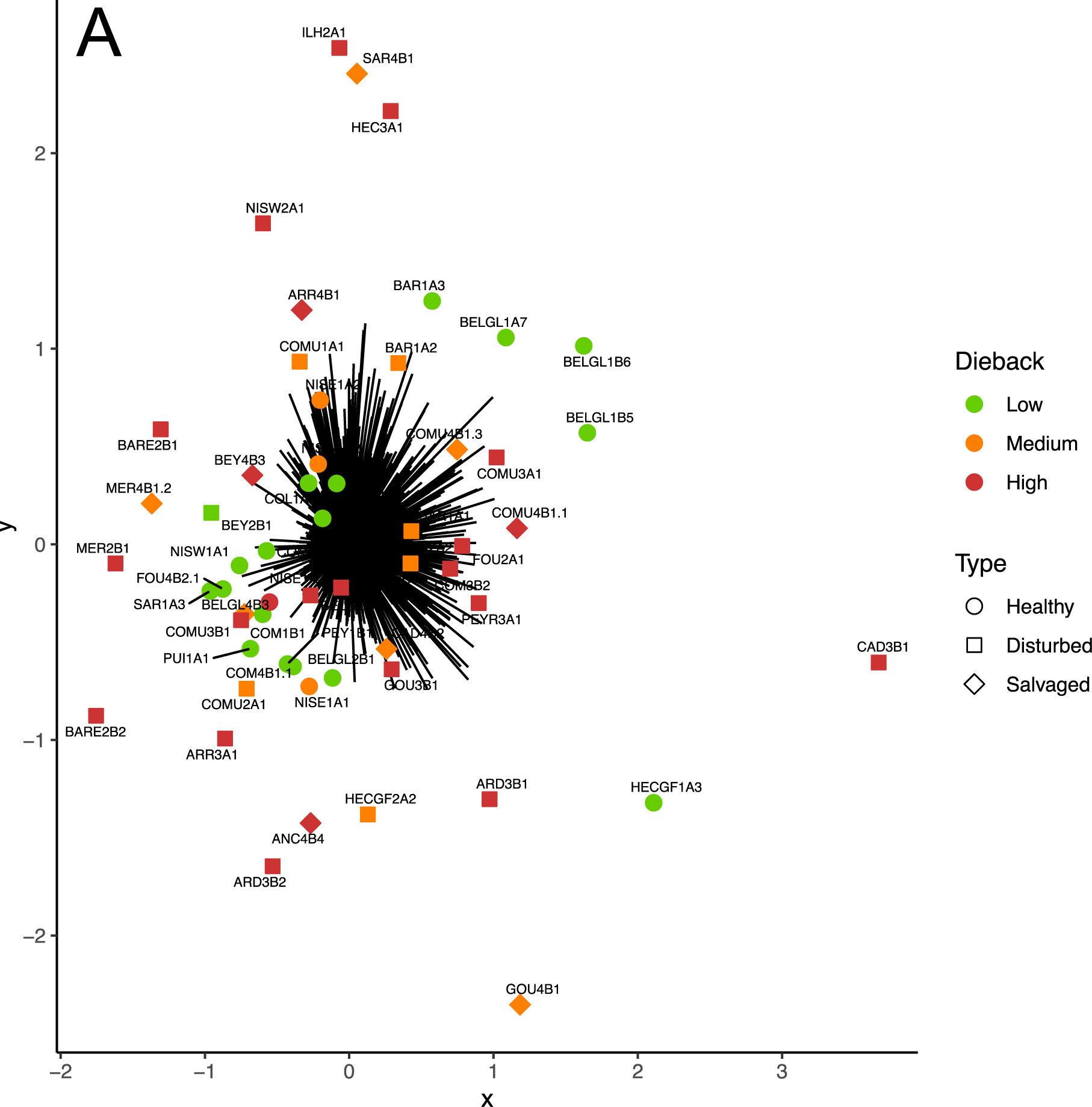
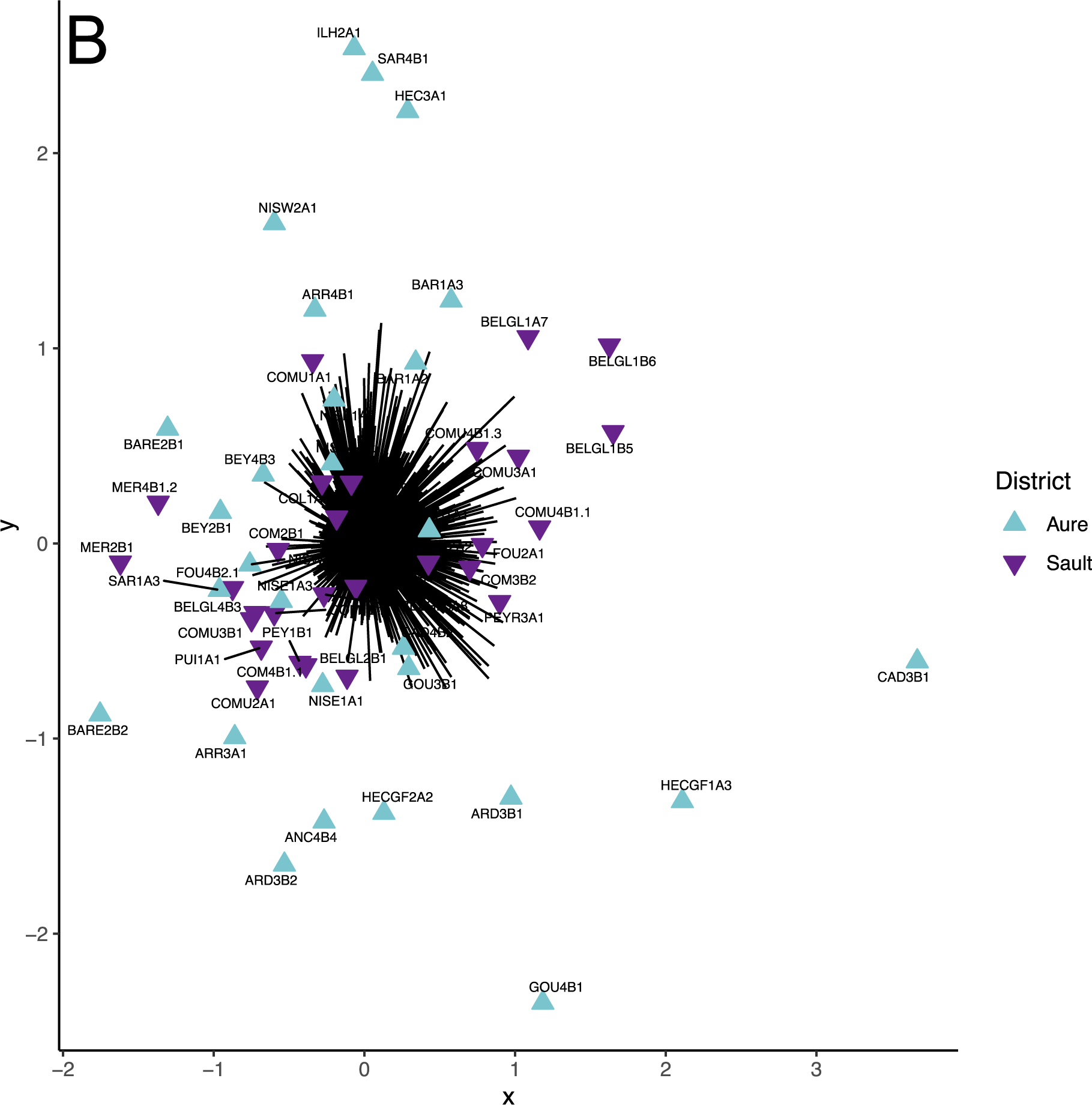
Variation in community composition across dieback gradient, stand types and geographical districts. Ordination plots representing variations in insect community assemblages in regards to: (**A**) dieback level conditions (*i.e.* low, medium and high) and stand types (*i.e.* healthy, disturbed and salvaged), and (**B**) to the two different geographical districts sampled. Mvabund analyses of community dissimilarities were performed using 999 bootstraps. Ordination (**A**) highlighted higher similarity in community composition between low dieback level plots than the other two dieback levels, more similar to each other but expressing high variability in community composition. No grouping of salvaged-logged plots in terms of community composition could be distinguished. Ordination (**B**) highlighted the greater dissimilarity in terms of community composition within plots of Aure valley compared with Sault plateau that hosted species assemblages more similar to each other.

### Winners and losers of forest disturbances

Zeta decline analyses allowed us to assess community assemblages and species retention rates across plots for the five main insect orders represented and for functional groups (floricolous / non-floricolous and parasitoids / non-parasitoid species) within each dieback category and stand type. We detected that species were retained differently within both dieback intensity gradient and stand types, regardless of the model scheme used (Fig. 5–6; Supplementary Fig. 6–7). Both Lepidoptera and Hemiptera showed a quick drop and a lack of structure in zeta ratio along the dieback gradient (Fig. 5A–C) and between the different stand types (Fig. 6A–C), indicating no species shared across zeta order ranges of both environmental gradients. Similar zeta declines were observed at higher zeta orders for Coleoptera at low dieback level and in healthy stands (Fig. 5A; Fig. 6A), as well as for Diptera and Hymenoptera at high dieback levels and in disturbed stands (Fig. 5C; Fig. 6B). These results highlighted that rare species shaped Coleoptera communities mostly at low dieback level, whereas common Coleoptera species could be found across plots of higher dieback gradients. Conversely, both Diptera and Hymenoptera species assemblages were less diverse at low dieback level and healthy stands while high dieback level and disturbed stands favoured complete species turnover with no common species retained (Fig. 5A–C, Fig. 6A–B). Regarding functional assemblages, all had common species likely to be found across low and medium dieback gradient, as well as healthy and salvaged forest stands (Fig. 5D–E; Fig. 6D–F). Nevertheless, while a similar pattern was observed for non floricolous and non-parasitoid species assemblages at high dieback level or within disturbed stands, zeta ratio of decline for both parasitoids and floricolous species cohorts fully dropped, indicating species compositional turnover within the two functional groups and no core species shared across 18 to 23 zeta order range (Fig. 5F; Fig. 6E). This lack of structure was likely driven by effects of high dieback level and disturbed stands observed on both Diptera and Hymenoptera, which included many taxa of pollinators and/or parasitoids (Fig. 5C, Fig. 6B). Overall, while few drops in species retention rates due to geographic effect were noticeable throughout the different environmental conditions (Fig. 5–6), main effects of both dieback level and stand types on species retention rates remained visible in ALL model scheme (Supplementary Fig. 6–7).

**Fig. 5:**
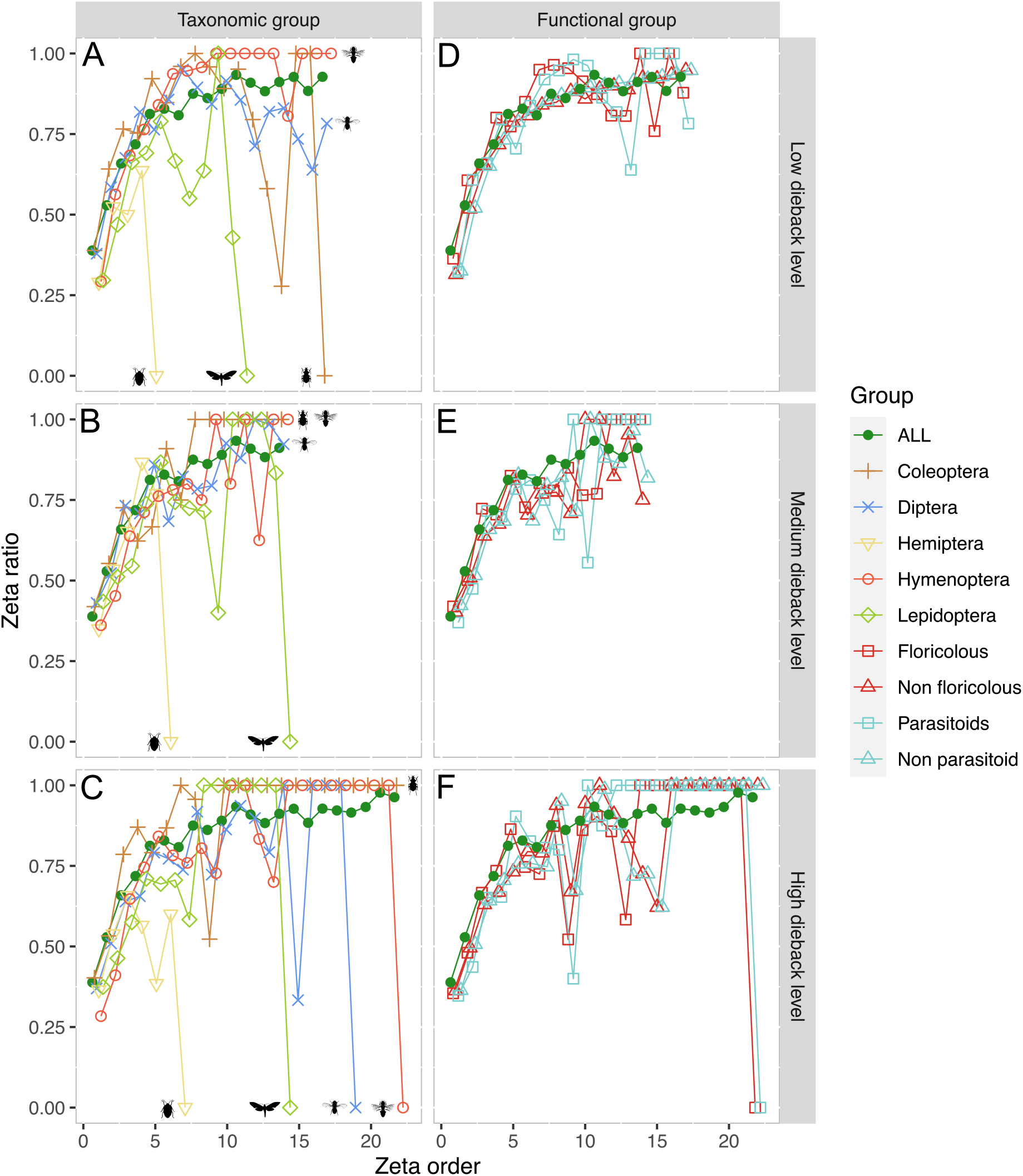
Effect of dieback on community composition for insect orders and functions. Representation of the species retention rate (*i.e.* zeta ratio) per plot (*i.e.* zeta order) following nearest-neighbour plot combinations scheme (NN) for low, medium and high dieback levels, respectively (**A, B, C**) for the five main insect Orders (Coleoptera, Diptera, Hemiptera, Hymenoptera and Lepidoptera) and (**D, E, F**) for the four main ecological functions recovered from taxonomic assignment (floricolous / non-floricolous and parasitoid / non-parasitoid species). Green line with plain dots represents mean species retention rate of the total dataset in each respective dieback category. Increasing curves express that common MOTUs are more likely to be retained in additional samples than rare ones (with presence of common species over all plots if zeta ratio = 1) and decreasing curves indicates species turnover. In our study, no core of common species could be sampled for Coleoptera at low dieback level as well as for Lepidoptera and Hemiptera throughout the entire dataset for each dieback category, respectively. For both Diptera and Hymenoptera, common core of species was observed within all dieback level plots except between high dieback level ones. Similarly, drop and lack of structure in common species was detected for floricolous and parasitoid functional assemblages within high dieback level plots while stable for the other two dieback levels, with this functional turnover being driven by dipteran and hymenopteran species turnovers at high dieback.

**Fig. 6:**
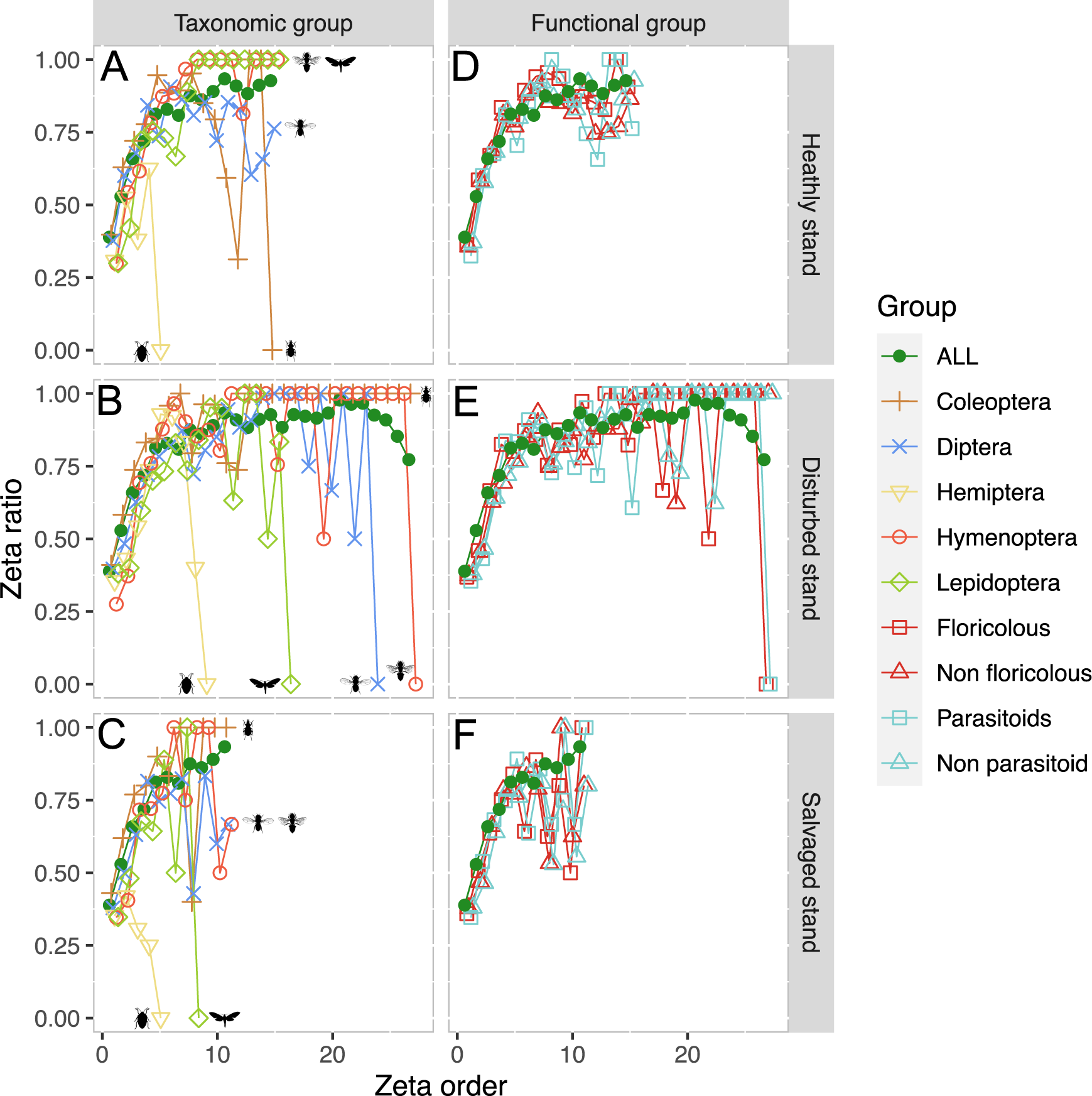
Effect of stand type on community composition for insect orders and functions. Representation of the species retention rate (*i.e.* zeta ratio) per plot (*i.e.* zeta order) following nearest-neighbour plot combinations scheme (NN) for healthy, disturbed and salvaged stands, respectively (**A, B, C**) for the five main insect Orders (Coleoptera, Diptera, Hemiptera, Hymenoptera and Lepidoptera) and (**D, E, F**) for the four main ecological functions recovered from taxonomic assignment (floricolous / non-floricolous and parasitoid / non-parasitoid species). Green line with plain dots represents mean species retention rate of the total dataset in each respective dieback category. Increasing curves express that common MOTUs are more likely to be retained in additional samples than rare ones (with presence of common species over all plots if zeta ratio = 1) and decreasing curves indicates species turnover. In the present case, we found no core of common species for Hemiptera within each studied stand type. For clarity purposes, Hemiptera are hereafter not considered. Within healthy stands, only Coleoptera had no common species retained, while Diptera had a slight drop in the probability of common species occurring, but yet not a complete compositional turnover. Lepidoptera, Diptera and Hymenoptera were all impacted by disturbed stands with complete compositional turnovers within studied plots, and similar observation could be noticed for parasitoid and floricolous functional assemblages. Meanwhile Coleoptera had a constant core of common species across the considered plots, and similar observation could be made within salvaged plots. Again, Lepidoptera assemblages were fully heterogenous across salvaged logged plots, and both Diptera and Hymenoptera expressed a lower probability of sampling common species, but yet not showing a complete change in community composition.

**Fig. 7:**
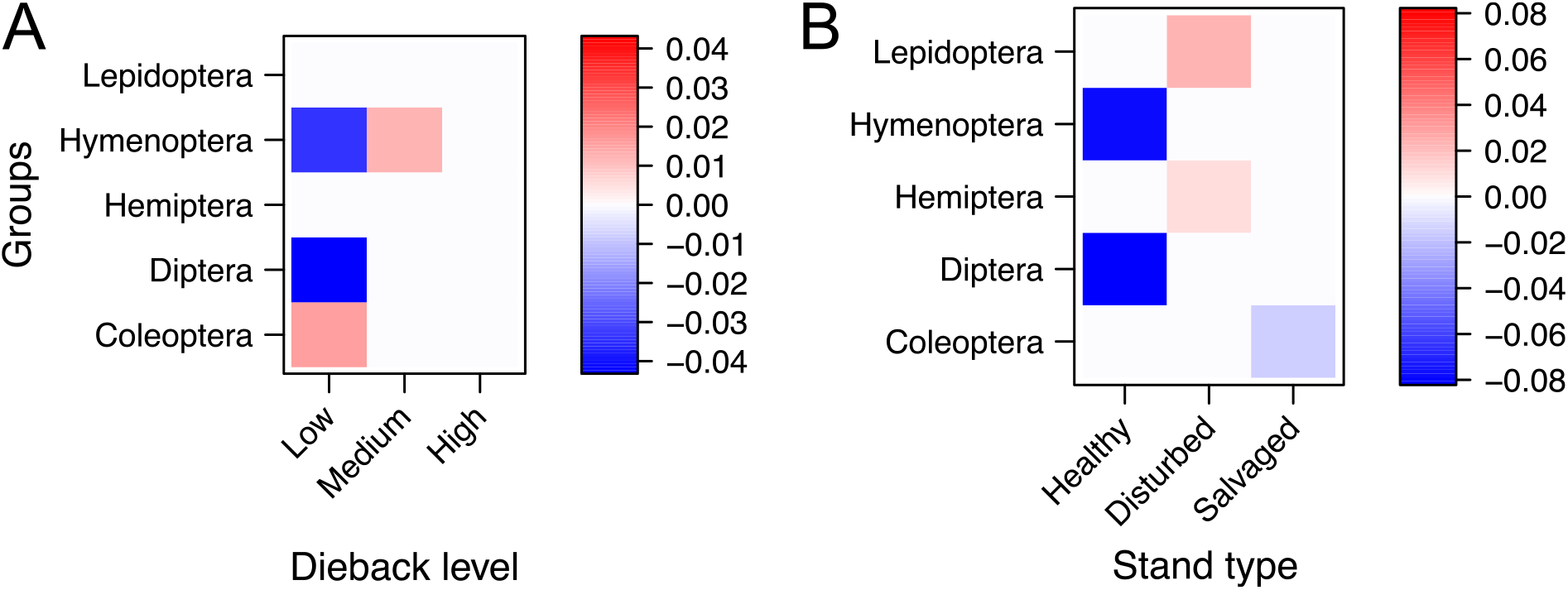
Insect orders winning and losing from environmental conditions. Heatmap representation based from traitGLM analyses of insect orders’ prevalence according to environmental conditions: (**A**) forest dieback level (*i.e.* low, medium and high) and (**B**) stand types (*i.e.* healthy, disturbed and salvaged). Negative and positive interactions and their intensity in terms of biodiversity response to environmental conditions for each of the five most represented orders are highlighted by a continuous colour gradient spanning from blue to red, respectively. We found specific responses for each insect orders’ diversity in regards to dieback levels and stand types. For instance, the low prevalence of dipterans and hymenopterans observed at low dieback level while relatively high prevalence for coleopterans. However, hymenopteran prevalence was greater in forests of medium dieback level. Coleoptera was the only order with prevalence negatively impacted by salvage logging. Finally, disturbed plots showed a relatively high prevalence of both Lepidoptera and Hemiptera.

Winning and losing insect orders in terms of prevalence over dieback gradient and stand types were assessed using a “fourth corner” modelling. Thus, we highlighted a higher prevalence in low dieback level stands and conversely, a detrimental effect of salvage logging over Coleoptera (Fig. 7). Furthermore, the lack of drop in species retention rate for both Hymenoptera and Diptera at low dieback level or in healthy stands (Fig. 5A, Fig. 6A) could be explained by a lower prevalence of these two orders in these environments (Fig. 7). Interestingly, Hymenoptera but not Diptera were winning from a particular level of dieback, especially with a higher prevalence at medium dieback level (Fig. 7A), while both Lepidoptera and Hemiptera diversity were favoured by stand disturbances in general (Fig. 7B).

Finally, by analysing congruences between MOTUs and dieback gradients using IndVal analyses, we highlighted species-specific responses to forest dieback. We significantly associated MOTUs to low and medium dieback level categories but not to high dieback level one (Supplementary Table III). These MOTUs could be linked to specific forest dieback conditions with particular environmental niches and therefore be considered losing over the general dieback gradient. The remaining species may either be too poorly sampled across plots to assess significant linkage or likely to be spread across multiple levels of dieback.

### Environmental drivers of species turnover

When assessing the contribution of our eight variables as drivers of the compositional turnover across zeta orders, we found that distance between plots played an important role in explaining the observed variance, especially in a two-plots to 10-plots comparison—hence dominated by rare species—and even greater at zeta orders above 28, thus on common species (Fig. 8A). This increase in geographic effect was in accordance with the relative number of plots in each respective district, for which communities were significantly different (Fig. 4B; Supplementary Table II). Besides distance, both altitude and canopy openness, and to a lesser extent density of large trees played a major role in driving community composition of rare species (Fig. 8A, zeta order 2). Similar results were observed at zeta order 10, but TreM diversity and volume of deadwood became more important, while canopy openness was less impactful. Finally, the bigger the zeta order was, the more impact had TreM diversity and deadwood volume on-site, as only these two environmental factors (distance excluded) were found to drive compositional turnover (Fig. 8A, zeta order 50). Interestingly, TreMs density did not affect overall compositional turnover (data not shown), suggesting that TreM diversity rather than TreM amount drove community composition. The eight tested variables explained 20 to 40% of the species composition turnover variance across all zeta orders. Distance excluded, the seven remaining variables only accounted for 15 to 20% of the variation, hence most of the variance remained unexplained (Fig. 8B). This suggests a complex multifactorial effect of environmental factors—but not random assemblies (Fig. 3C–D; Supplementary Fig. 5C–D)—, with many variables yet to be explored, in driving the insect community composition turnover of both rare and common species.

**Fig. 8:**
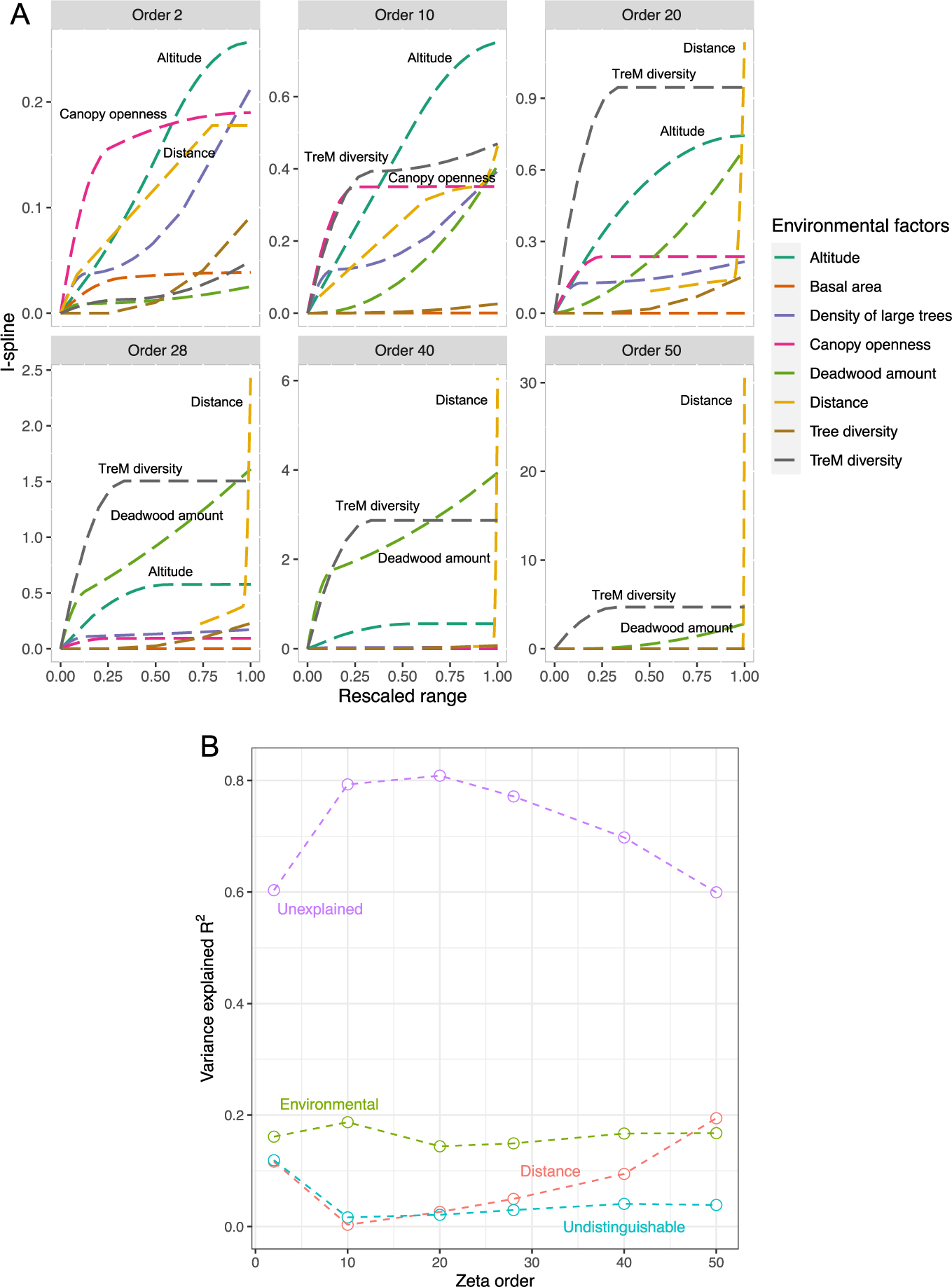
Environmental variables driving species community changes (zeta diversity changes). (**A**) I-splines values of predictors for seven environmental variables with their respective contributions to zeta diversity. Environmental contributions are shown for six different zeta orders (*i.e.* ζi) here referring to plots compared together (from ζ2 to ζ50), with the main contributively environmental variables highlighted for each zeta order. We observed a shift in the key forest features explaining community changes according to geographic scale, with canopy openness and altitude as main environmental drivers at local scale (ζ2), replaced by tree-related microhabitats and deadwood amount at large scales (from ζ28 to ζ50). (**B**) Proportion (in r^2^) of the total zeta diversity variance explained by environmental variables, distance, undistinguishable variables or unexplained, according to zeta order. The seven tested environmental variables accounted for 15–20% or the total variance in community changes regardless of zeta order, with distance explaining up to 20% of the total variance as well depending on the different zeta orders considered.

## Discussion

Marked declines in insect abundance, biomass and species richness have recently been quantified in Europe^2^. The causes of insect decline are multifactorial with rapid climate change identified as one of the major drivers. Here we investigated whether forest disturbances such as drought-induced forest decline and subsequent salvage logging have an impact on flying insects. Surprisingly, insect richness remained stable regardless of the extent of dieback or logging. However, species composition changed significantly with the level of dieback, driven mainly by rare species. Comparable changes in insect species composition but stable insect species richness have been observed in response to pest-induced forest dieback of other tree species^22, 33^, and the fact that similar observations have been made in aquatic insect communities over time^17^ suggests that our findings are indicative of broad patterns.

While species richness is similar, we highlighted that forest dieback caused significant changes in insect community compositions for the whole dataset. In an increasing impact order of dieback level, Lepidoptera, Hemiptera, Diptera and Hymenoptera all showed changes in community composition across the dieback gradient. Furthermore, all functional groups studied (*i.e.* floricolous / non-floricolous and parasitoids / non-parasitoids) showed compositional turnover driven by forest dieback. In line with previous study, we demonstrate how widely impacted biodiversity can be from dieback regarding both taxonomic and functional assemblages^34^. Only Coleoptera have had no detected effect of disturbances in general on their communities. This is surprising as Coleoptera have been shown to be significantly affected by both forest diebacks and salvage logging^35^. But while they were the sole group bearing significant change in species richness across districts, this absence of community change across dieback gradient had been already observed on saproxylic beetles in the same sampling area^36^ but may also be an artefact of the use of Malaise traps and a biased or lower sampling efficiency of Coleoptera in Sault plateau.

Salvage logging practices had no impact on species richness nor on community composition in the Pyrenees. This result has to be taken with caution since the salvage logging intensity and the associated amount of deadwood removed from our plots might have been too small to significantly impact insect fauna. Indeed, as the slopes of the sampled mountain forest plots were steep and logging mechanically performed using skidders and cables, we suppose that both reasoned management practices and the difficulty to heavily harvest deadwood mitigated salvage logging impacts on insect biodiversity. In addition, winning and losing taxa from salvage logging^13^ may have had overall counter-balancing effects. Indeed, we found a weak but expected negative effect of salvage logging on the prevalence of Coleoptera similar to other studies^13^. However, we think the first explanation of low salvage logging intensity is more plausible and supported by the fact that we did not detect any impact of salvage logging on deadwood amount in our study plots^36^. Hence, one must remain cautious with the lack of detectable impact of salvage logging in our study. Furthermore, as sampling was happening in non-old-growth forests, this may have left a sufficient amount of coarse woody debris on-site to prevent severe impacts on insects. Insect communities in managed stands such as those we studied may be poorer and more homogenous than in more mature forests, reducing our ability to detect changes in species richness. Additionally, our sampling was carried out more than 10 years after the onset of climate-induced tree dieback and subsequent logging, giving plenty of time for a recovery of species richness, even though extinction debt (*i.e.* a time-delayed negative response of a taxon to an environmental disturbance) has already been observed in Diptera after 29 years in *Quercus* spp. dominated forests of Southern France^37^.

Overall, our findings underline that forest dieback favouring insect biodiversity^14^ may be based on the response of well studied groups, *i.e.* “those insects that we love and cherish”^38^. Indeed, rather than an increase in biodiversity, we demonstrated that species richness remained steady regardless of dieback level. Even though they did find an overall positive response, Moretti *et al.*^15^ highlighted by breaking down the datasets at lower taxonomic level that insect diversity could respond both negatively and positively to fire-induced forest diebacks. They found that the winners were among the most studied groups of insects (*i.e.* hoverflies, bees, social wasps and ground beetles) while only one losing insect group was identified (weevils)^15^. This implies that other, unidentified groups were potentially “losers” but less likely to be reported. Similar bias toward these well-known insect groups is also identifiable in studies on pest-induced forest dieback with an additional focus on red-listed species^10^, or in the studies considered in Thom and Seidl’s meta-analysis^14^. From this detailed taxonomic view^15^ and recent comprehensive studies^22, 33^, insect positive response to disturbances must be taken cautiously as it could be biased towards the response of the better studied insect groups only. Previously reported global patterns may thus not be fully representative. Here, by including hyper-diverse and understudied groups (“dark taxa”)^32^, the overall stable species richness that we found across both the dieback gradient and stand types highlights a nuanced and idiosyncratic response of insect orders and functional groups such as floricolous / non-floricolous and parasitoid / non-parasitoids insects to forest disturbances. Additionally, our results in both insect species richness and compositional turnover emphasize the limitation inherent to the sole use of species richness as a metric^39^ and the need to look more closely at community composition and functional changes to notice disturbance effects^40^.

Here, we highlight that various dieback levels would rather promote different insect community assemblages and be part of natural succession dynamics (*i.e.* ecological change occurring in a predictable way after disturbance), with equally weighted response of species either winning or losing throughout the dieback intensity gradient. Hence, management policies based on species richness metrics and promoting dieback only to favour biodiversity may hold potential risks by driving a catastrophic decline of insects through a potential homogenization of dieback levels and associated species assemblages with more generalist species. Albeit Borges and collaborators^40^ highlighted the issue in the very distinct island ecosystem, their concern can be extended to freshwater streams^17^ or disturbed forest environments as in the present case.

In our study, zeta decline fitting a power-law regression implied a compositional turnover of the total dataset driven by environmental factors rather than stochastic events^29^. The zeta ratio analyses showed that species are indeed retained differently within each dieback category. Thus, the lack of core set of species observed for Lepidoptera may be attributed to a relatively low moth diversity associated with silver fir-dominated forests with community composition mainly shaped by turnover processes^41^. On the other hand, sampling efficiency could explain the absence of common species retained for Hemiptera for the different dieback categories or stand types tested^29^. Nevertheless, the prevalence (*i.e.* the presence probability of a taxa within all the plots grouped in a given environmental category) of these two orders remained positively correlated with dieback in general, in accordance with previous observation in which open canopy deriving from diebacks favours heliophilous and flower-visiting insects^15^. Interestingly, the prevalence of Coleoptera was favoured by low dieback level while species richness across the dieback gradient was similar and the species retention rate decreased. Even though we could not finely assess feeding guilds, these observations may support high turnover processes and competition deriving from specialist species’ population decrease as factors shaping Coleoptera communities^9^. However, salvage logging had a slight but significant negative impact on Coleoptera prevalence, in line with previous observations^13, 36, 42^. In addition, each dieback gradient hosted a specific core set of one or more common species composed of Diptera and Hymenoptera at both low and medium dieback levels. The reduced openness associated with healthy stands—hence low dieback level—might also support the reduced prevalence of both Diptera and Hymenoptera observed in these environments, as most of the species recovered for these orders are considered floricolous. Dieback might also increase environment complexity and support the positive effect of medium dieback level that we observed on Hymenoptera, especially parasitoid ones^23^, rather than a negative bottom-up effect of the host condition on parasitoid species^24^.

A compelling observation in terms of functional diversity is that while both Diptera and Hymenoptera displayed strong compositional turnover at high dieback level, their drastic changes in community composition drove the turnover observed for floricolous and parasitoids functional assemblages. It is known that both floricolous and parasitic Hymenoptera can indeed be significantly impacted by structural changes in silver fir-dominated mountain forests^43^. However, our results emphasize the ecological importance of some species-rich yet often overlooked taxa on wide functional groups. Furthermore, it highlights the need for more work to investigate whether climate-induced forest dieback can deeply impact functional efficiency and ecosystem services through these taxa^3^.

In a more detailed view, within each studied order or functional group, the winner and loser taxa were relatively rarely occurring species in our dataset. This pattern is comparable to previously reported taxonomic dissimilarities driven by rare species^34^ and consistent with models showing that selective perturbations directed towards rare species generate more dynamical effects on the ecosystem^44^. As rare species have a positive impact on ecosystem multifunctionality^45, 46^, these rare taxa have high patrimonial value and thus their importance in forest ecosystems functioning should be further investigated^34^.

Geographic distance—regardless of the zeta order considered—played a significant role as a macroecological factor shaping the different insect communities of the two districts and, as for the aforementioned winner and loser taxa, mostly taxa with low prevalence shaped compositional specificities of each district. As expected, many forest features tested (*i.e.* altitude, density of large trees, canopy openness, total volume of deadwood and TreM) also had a significant effect in shaping insect community composition, indicating a complex response to disturbances. Remarkably though, environmental factors explaining most of the compositional variance differed according to the observed zeta ratio. At low scale (zeta order = 2), community changes were mainly driven by canopy openness, density of large trees and altitude. This result support the role of distance and stand structure heterogeneity in favour of biodiversity^47^ and is in line with previous observations on these features affecting the rare species composition at low zeta order^22^. However, these environmental features at large scale (zeta order > 10) were no longer the main drivers of biodiversity changes. Instead, both TreM diversity and the amount of deadwood were the main factors influencing community composition when considering the whole dataset. Interestingly, TreM diversity rather than density had most impact on insect diversity (data not shown)^48^, similarly to deadwood diversity and amount in other studies^47, 49^. However, as TreM diversity is linked to TreM density, it therefore might indirectly and at different intensity also influence biodiversity^48^. Furthermore, and contrary to previous report^47^, we found deadwood amount to be as valuable as stand structure for biodiversity conservation, depending on the geographic scale managed^42^. This result further supports the general call on acknowledging and considering the importance of deadwood to biodiversity^42^. Interestingly, a previous study successfully linked hoverfly diversity to plot connectivity and the combined effect of environmental factors from both stand and landscape scales^37^. The shift in environmental variables driving community assemblages according to the zeta order considered we highlighted here support such geographic scale effect and help disentangling the key drivers in action. This may potentially be extended to most aerial insects with higher dispersal ability, which are the main representatives of our Malaise-trapped dataset. Hence, while species-specific response to environmental factors must be accounted for, geographic scale of the conservation area, especially through stand and landscape scales^37^, as well as dieback intensity should also be considered when managing stand heterogeneity and environmental factors to favour forest insect biodiversity^47^.

Finally, our study confirms DNA metabarcoding of Malaise trapped samples as an efficient approach for biomonitoring changes in species-rich insect communities that include “dark taxa” which collectively represent the bulk of forest insect diversity^22^. Indeed, our metabarcoding pipeline coupled with recent efforts to complete the DNA barcoding reference library of European insect fauna^19, 50^ allowed the inventory of nearly 3000 insect species, more than half of which were identified to species level, including hyper-diverse taxa such as Diptera and Hymenoptera^51^. These two taxonomic groups were the most diverse in our study, like in other Malaise trap environmental surveys^52, 53^. Although Malaise-trapped samples are biased toward flying insects^54^, the zeta decline analyses indicated that our dataset incorporated rare species as non-stochastic events and tracked changes in both species richness and community composition at both large and small spatial scales while overcoming sampling district discrepancies^29^. Scaling up our approach to national and continental levels would help to monitor insect biodiversity and potential decline, to provide an understanding of the environmental drivers of biodiversity loss in a rapidly changing climate, and allow for regular assessment of the efficiency of conservation and forest management policies.

## Methods

### Study sites, stand description and insect sampling

Fifty-six plots were set up in 2017 in silver fir forests in Central (Aure valley) and Eastern (Sault plateau) French Pyrenees (see “*Supplementary Information – Plot list*”).

Tree dieback was assessed using the ARCHI method^55^. The 20 closest silver firs from the plot center (defined by the Malaise trap), belonging to the dominant layer and with a diameter at breast height (dbh) > 17.5 cm, were examined with binoculars. The proportion of affected trees (*i.e.* trees expressing signs of climate-induced dieback and comprising stressed + resilient trees) and dying trees was used as the proxy of the decline at the stand level to define a three-category gradient: 18 low dieback level plots (no dying trees and ≤ 60% of affected trees), 23 high dieback level plots (at least 1 dying tree and > 60% of affected trees) and 15 medium dieback level plots (no dying trees and > 60% of affected trees).

For each living or dead tree belonging to a fixed-angle plot (ratio 1/50), we recorded its status (living, snag, log), tree-species, dbh (minimum dbh to be recorded: 17.5 cm for living trees and logs, 7.5 for snags). Tree-related microhabitats (TreMs) were recorded on living trees and snags according to the Larrieu and collaborators’ typology (47 types)^48^. TreM diversity corresponds to the number of TreM types recorded per plot. Very large trees were defined as living trees above 67.5 cm in dbh. Since we used the ratio 1/50 to fix the angle, basal area matches with the number of trees observed. Each deadwood item (length > 1 m) was measured in length and diameter to calculate its volume. Mean plot area was about 0.3 ha, depending on the dbh and location of the largest trees. Additionally, canopy openness was evaluated by using a spherical densiometer^56^ at the plot centre and 10 m from the centre in four crossed directions (thus 5 locations per plot). Canopy cover was calculated for each location as the proportion of the 68 points of the densiometer screen that was intersected by cover. Plot value resulted as the mean of the five counting.

Stand type was described on an area of 1 ha using the Index of Biodiversity Potential (IBP)^57^. The IBP is a biodiversity evaluation tool (according to Larsson^58^) combining ten historical, structural and compositional key factors for forest-dwelling species which are easily and directly measurable in the field^59^. The measurement of these factors provides a rapid assessment of the theoretical biodiversity hosting capacity of the stand and facilitate the identification of the key environmental variables manageable for better biodiversity potential. In our study, the main factors accounting for healthy, disturbed and salvaged conditions were stand structure (canopy openness), tree health (deadwood volume and number of living trees) and the awareness of management activities influencing deadwood volume and density of habitat-trees available on-site.

Insect sampling was conducted over late spring–early autumn season from May 15 to September 15, 2017 using 56 Townes-style Malaise traps with black walls and white roof^53^. One Malaise trap was placed at the centre of each of the 56 one-ha plots. Malaise trap samples were filled with a mixture of 20% mono-propylene glycol and 80% pure ethanol. Samples were retrieved once a month, giving a total of four samples per plot over the sampling period (124 trap days), for a total of 224 samples. After collection, all samples were stored at 4°C until laboratory processing.

### Laboratory processing and DNA extraction

Insects were first filtered from the trapping solution and rinsed with ultrapure Milli-Q water to remove mono-propylene glycol residue. Insects were then placed within sterile and disposable Petri dishes on clean absorbing paper to dry overnight at ambient temperature.

Once dried, insects were size-sorted using decontaminated forceps. Insects larger than a European honey bee (*Apis mellifera* Linnaeus, 1758) were removed and only the head or a part of the abdomen was retained in order to reduce the biomass and improve the detection of rare or small species^60^. Insect bulk samples were then ground and homogenized into fine powder using disposable BMT-50-S-M gamma sterile tubes (IKA) with 10 steel beads with an Ultra Turrax Tube Drive grinder (IKA). Ground bulk samples were then conserved at -21°C.

DNA extraction of 25 mg (±2 mg) of insect powder was performed on silica columns using a standard DNeasy® Blood & Tissue extraction kit (QIAGEN) (see: www.qiagen.com/handbooks for further information). For each ground sample, we took 25 mg (±2 mg) of insect powder in a 1.5-mL microcentrifuge tube using spatula previously decontaminated with 4% Decon® 90 solution and autoclaved. During powder sampling, empty 1.5-mL microcentrifuge tubes containing 200 µL ATL buffer were left open and changed every nine samples to control for potential cross-contamination with volatile insect powder and all were processed as extraction controls (EC). Mock communities of 248 Asian insects were used as positive controls. Negative (NC), extraction (EC) and positive (PC) controls were processed down to sequencing. Lysis was performed with horizontal shaking overnight at 56°C in 180µL ATL buffer and 20 µL proteinase K. All vortex steps were replaced by handshaking to reduce DNA degradation and DNA was eluted into 80 µL of AE buffer following 15 min incubation on the silica column at ambient temperature, and a second elution with the previous eluate after 5 min incubation on the silica column. Each sample was quantified using Qubit® 2.0 fluorometer dsDNA High Sensitivity kit (Invitrogen) and eluate subsamples were diluted to 2 ng/µL. Both stock solutions and diluted eluates were stored at -21°C.

A twin-tagging dual-indexing approach was used for the sequencing library preparation^61^ (see “*Supplementary Information – Primer list*”). We targeted a 313-base pair (bp) fragment of the mitochondrial DNA Cytochrome oxidase *c* subunit I (COI) gene. PCR amplification was done using the mlCOIintF forward primer 5’– GGWACWGGWTGAACWGTWTAYCCYCC–3’ and jgHCO2198 reverse primer 5’– TAIACYTCIGGRTGICCRAARAAYCA–3’^62, 63^. To facilitate twin-tag multiplexing, each of the 96 primer pairs was synthesized with a 7-bp tag at their 5’ end, differing by at least three bp and avoiding G nor TT at the 3’ end to prevent a succession of the three same nucleotides once attached to the primers. Furthermore, A, T, CA, CG, GC and GT nucleotides were added to the tag to act as heterogeneity spacers^64^ to increase complexity and shift the reading frame for better sequencing quality on Illumina MiSeq platform (see “*Supplementary Information – Primer list*” for a complete list of designed and used primers). Shifting was in accordance with green/red light balance on v2/v3 Illumina MySeq technology^65^. Neither proof-reading nor hot-start Taq polymerases were used for PCR amplification due to Inosin bases in the reverse primer^66^.

Prior to amplicon PCR amplification, we tested the optimal number of cycles and DNA quantity added to the PCR-mix by quantitative PCR (qPCR) using a LightCycler® 96 Instrument (Roche) with the KAPA Library Quantification Kit (Illumina). The qPCR conditions were: one preincubation step of 95°C for 5 min, followed by 40 cycles of 95°C for 30 sec, 45°C for 60 sec and 72°C for 90 sec, with a final extension at 72°C for 10 min and a high-resolution melting step of 95°C for 60 sec, 40°C for 60 sec, and an acquisition gradient from 65°C to 97°C. The qPCR reactions with a final volume of 15 µL were prepared with 7.5 µL of KAPA qPCR mix (2X), 0.3 µL of each forward and reverse primers (10 mM), 3.9 µL of extra pure molecular grade water and dilution series of 3 µL DNA template (set at different concentrations: 8, 4, 2, 0.4 and 0.2 ng/µL) were performed to investigate inhibitions. The optimal number of PCR cycles was designated by where the exponential phase transitioned to plateau phase. PCR amplifications were thus run for 23 cycles under the same conditions as the qPCR, in 25 µL reaction volume as follows: 2.5 µL of buffer green (10X), 0.5 µL of dNTPs (10 mM), 1 µL of each forward and reverse primers (10 mM), 0.2 µL of Metabion mi-Taq (5 U/µL), 16.8 µL of extra pure molecular grade water and 3 µL of DNA template at 2 ng/µL. Each sample was PCR-amplified three times in different 96-well plates with three different twin-tag sequences, to allow independent tracking of the three PCR products after amplification. PCR products were purified using CleanPCR magnetic beads (CleanNA) at a ratio of 0.8/1 µL of PCR product. Two steps of rinsing with 200 µL of 70% ethanol were performed before final elution in 25 µL TE buffer (1X). Purified PCR product was then quantified in duplicates on a FLUOstar OPTIMA microplate reader (BMG Labtech) with Quant-iT^TM^ PicoGreen® dsDNA assay kit (Thermofisher) following manufacturer’s protocol. Equimolar pooling of the samples was carried out for each plate with a total DNA quantity of 1 µg of purified amplicon for a final volume of 60 µL. A second indexing for all PCR plates was conducted using the TruSeq DNA PCR-free Library Prep kit for PCR-free ligation (Illumina) with 18 different indices (one per 96-well plate) following the protocol given in Leray *et al.* (2016)^61^. Library preparation and sequencing of the total 224 was done on five MiSeq runs – v3 (Illumina) with 600 cycles.

### Use of positive controls and blanks in demultiplexing

Mock communities of known species composition with DNA barcode sequences available were used as PCs. The 248 species used in the mock community all have distributions restricted to east Asia, and hence allow for the examination of tag-jumping in the Illumina sequencing process. PCs were sequenced and bioinformatically treated the same way as bulk samples to ensure comparability. We focused on a restrictive approach, favouring false negatives to false positives (See Alberdi *et al.,* 2018^67^ for more information on bioinformatic set-up). To decide both on the number of PCR replicates in which MOTUs must appear and minimum read numbers per PCR, we used BLAST+^68^ on the PCs to find the set up in which we could recover the most MOTUs with 100% match and no MOTUs below 97%. Both NC and EC were checked for reads after demultiplexing to ensure that there was no cross-contamination and removes potential MOTUs recovered in controls from the total dataset.

### Reads demultiplexing and taxonomic assignment

AdapterRemoval *ver. 2.2.2*^69^ was used to trim the twin-tagged adaptors of the reads and we employed sickle *ver. 1.33*^70^ to perform paired-end quality trimming. Error correction using Bayes Hammer was done via SPAdes^71^ *ver. 3.12.0,* followed by paired-end merging with PandaSeq^72^ *ver. 2.11*. PCR replicates were treated following DAMe pipeline^73^. Heatmaps of primer tag combinations versus read numbers were generated using R *ver. 3.6.1*^74^ to control for indexing quality. Our twin-tagging approach allowed us to easily identify errors and problems at this step. Renkonen Similarity Indices (RSI value^75^; were calculated for unique sequences found in at least 3 PCR replicates and represented by at least 3 reads (See section ‘<u>Use of positive controls and blanks in demultiplexing</u>’ for arbitration of required numbers of PCR replicates and reads). Average read size was plotted in R to ensure filtering and trimming quality, and VSEARCH^76^ *ver. 2.8.1* was run to remove chimeras.

Sequences were clustered into Molecular Operational Taxonomic Units (MOTUs) with a 97% similarity threshold^77^ using SUMACLUST *ver. 1.0.20*^78^. Clustering quality was controlled via R, using the LULU approach^79^.

Finally, taxonomic assignment was performed against BOLD System database^27^ in April 2019 using a 97% threshold similarity^22^ with bold R-package *ver. 0.9.0*^80^. For each MOTU, only unambiguous taxonomic assignments were kept, meaning that when multiple names at a given taxonomic rank were attributed, only common and unique taxonomic assignments at higher taxonomic levels were kept. In the case of multiple MOTUs with an identical and unambiguous species name assignment, all related MOTUs and their respective presence in sampled plots merged into a single species-specific MOTU. Functional traits were associated manually to MOTUs according to family level whenever possible (see “*Supplementary Information – Family functional traits*” for more details). Parasitoid and floricolous families were mainly assessed from^81–83^ and with the help of expert taxonomists.

### Statistical analyses

All statistical analyses were carried out in R *ver. 3.6.1*^74^. Read-sequencing data were transformed in incidence-based data to account for presence / absence of MOTUs given that abundance data from metabarcoding can be misleading for metazoans^84^.

We generated accumulation curves using *iNEXT* package *ver. 2.0.20*^85^ with incidence-based frequency dataset parameters and Hill number *q* value set to 0 to reflect truly observed MOTU diversity, regarded as species proxy. Total number of MOTUs was plotted against both sampling units and sample coverage for the late spring–early autumn sampling period as well as for each four-weeks time series (named May, June, July and August from their respective starting months). MOTUs belonging to the five most diverse taxonomic orders represented in our dataset (*i.e.* Coleoptera, Diptera, Hemiptera, Hymenoptera and Lepidoptera) as well as for the remaining 10 Orders (*i.e.* Blattodea, Ephemeroptera, Mecoptera, Neuroptera, Orthoptera, Plecoptera, Psocodea, Raphidioptera, Thysanoptera and Trichoptera) pooled into an “Others” category, were also plotted against sampling units to account for respective recovery success.

We also used both Chao and first order Jackknife (Jack1) methods to have different biodiversity estimates from the ‘specpool’ function of R package *vegan ver. 2.5-6*^86^. These estimators also extrapolate species richness within the sampled area in regards to sampling effort and based on incidence data.

Impacts of tree diebacks on species richness was assessed using generalised linear model (GLM). For the total dataset, each taxonomic order category of the five main representatives (*i.e.* Coleoptera, Diptera, Hemiptera, Hymenoptera and Lepidoptera) or over the four ecological groups (*i.e.* floricolous / non floricolous adults and parasitoid / non-parasitoid larval feeding guild), GLM was first tested with both Poisson and quasi-Poisson distribution, the latter always retained for its better fit. Then, each model tested the effect of the dieback gradient, stand types, their interaction as well as the effect of districts. When no interaction was found, a simpler additive model was tested. In all nine studied cases, quasi-Poisson distribution and additive model characteristics were retained, as follow: `model <-glm(richness ∼ dieback + salvage + district, family = quasipoisson)`. We finally applied type II Anova from *car* package *ver. 3.0-8*^87^ on the model to test the significance of each parameter on the species richness of the studied category.

To compare insect community composition across disturbances gradients (*i.e.* tree dieback intensity and stand types) and districts, we performed GLM on a generated mvabund object using *mvabund ver. 4.3.1*^88^ as follow: `Mod_Tot <-manyglm(Tot_mvabund ∼ X$Dieback + X$Type + X$District, family=“binomial”)`. Reported values are the two extremes resulting from a 10 runs range. Post-hoc Holm correction was then applied on the 10 p-values recovered to assess for the significance of potential compositional changes in community compositions. Ordinations for these GLM were performed using *ecoCopula ver. 1.0.1*^89^.

Prevalence of the five most represented taxonomic orders in regards to dieback levels and stand types was assessed by a “fourth-corner model” analysis using ‘traitglm’ function from *mvabund ver. 4.3.1*^88^ R package. For each insect MOTU, taxonomic order was defined as a trait and GLM analyses run to define traits significantly associated to environmental covariates.

To determine potential indicator species for each dieback gradient, we used IndVal analysis from ‘multipatt’ function of *indicspecies* R package *ver. 1.7.9*^90^. Again, reported p-value ranges are resulting from 10 independent IndVal runs with post-hoc Holm corrections subsequently applied.

We also assessed the importance of rare species in species assemblages specific to both the gradient of dieback intensities and the sampling districts. Here, we defined a rare species from its frequency of appearance in a given dieback category or district, with occurrence (*i.e.* number of plots of the relative category in which we found the species) as an index of relative abundance. Hence, MOTUs more frequently sampled throughout the plots were defined as more abundant and more common. We then generated heatmaps of relative abundance using the phyloseq *ver. 1.30.0*^91^ package for R. As rare species—defined from the multiple dimensions of rarity^92^—can therefore be virtually locally abundant in a given dieback category or district by the use of occurrence only to define our MOTU rarity, relative abundance used in heatmaps was generated from 100 randomized bootstraps of the total dataset to virtually increase sampling effort. Bootstrapped plots were then concatenated to recover a bootstrapped score giving the relative abundance. From this score, we could infer rarity in terms of habitat specificity, with common species occurring across spatial scale and/or dieback gradient, and rare species occurring only in a few plots in a particular district or level of dieback. Each of the three dieback categories (*i.e.* low, medium and high dieback levels) was rescaled to account for the difference in sample representativeness, as such that each category was depicting one third of the total dataset. Each MOTU relative abundance was then calculated in percentage from the bootstrapped score for these rescaled categories, so that a MOTU present in all plots of one particular dieback category was present at 33% in the respective category, for a total of 100% across the total dataset if appearing everywhere. Regarding abundance per district (*i.e.* Aure valley and Sault plateau, as well as total dataset), MOTUs’ relative abundances were calculated in percentage based on the bootstrapped score with no further rescaling per district.

Species compositional changes along the total sampling area was assessed by calculating the number of species shared across all 56 experimental plots using zeta diversity (*i.e.* extension of beta diversity over *i* zeta order (ζi) or plots)^28^ using zetadiv R package *ver. 1.2.0*^30^. Two models were computed to perform plot comparisons either following a nearest-neighbour (NN) scheme or considering every comparison possible (ALL scheme) to assess for heterogeneity driven by distance. The two models (NN and ALL) were also fitted both to power-law (*pl*) and exponential (*exp*) regressions to define whether the signal in our data set was driven more by ecological niche and rare species turnover rather than by stochasticity, respectively^29^. To assess the better fit to a model between *pl* and *exp*, we chose the lowest score from Akaike Information Criterion (AIC) which imposes penalty to each model in regards to its number of parameters, and thus favour models satisfying parsimony criterion^93^. For both dieback gradient (*i.e.* low, medium and high dieback categories) and stand types (*i.e.* healthy, disturbed and salvaged), zeta ratio (*i.e.* the species retention rate, giving the probability of a common species to be retained in the community across zeta order here referring to plots) was generated for each of the five most represented taxonomic orders and the four different functional groups identified, as well as for the mean zeta ratio of the respective dieback category or stand type on the total dataset.

Estimations of environmental variables contribution to insect species assemblage changes were performed with zeta multi-site generalized dissimilarity modelling (zeta.msgdm) in zetadiv R package *ver. 1.2.0*^30^ at different zeta order, ranging from two to 50. In total, eight environmental variables were tested with zeta.msgdm: geographic distance to the nearest plot, altitude, canopy openness, total amount of deadwood in m^3^ per ha, the basal area per ha, the tree diversity per ha, the density of very large trees (Ø > 67.5 cm) per ha and the TreM diversity per ha. zeta.msgdm was calculated using I-spline models^94^.

## Supporting information

Supplementary information - Family Functional Traits List

Supplementary information - Plot List

Supplementary information - Primer List

Supplementary Table III

## Acknowledgements

We are grateful to all forest stakeholders—owners and managers—that allowed us to perform fieldwork in their stands. We are thankful to Laurent Burnel, Jérôme Molina, Sylvie Ladet, Carl Moliard, Jérôme Willm, Wilfried Heintz, Denis Sabadie and Benoit Nusillard for the field work, field and GIS support, as well as to Kévin Speder for his help in the molecular lab. We are also thankful to Yohann Graux for his help with QGIS mapping and Rodolphe Rougerie for his valuable comments on the manuscript. Library preparation for sequencing as well as sequencing on Illumina MiSeq platform were performed at the Berlin Center for Genomics in Biodiversity Research (BeGenDiv)—Königin-Luise-Straße 6–8, 14195 Berlin, Germany. This research is part of the international project CLIMTREE “Ecological and Socioeconomic Impacts of Climate-Induced Tree Dieback in Highland Forests” within the Belmont Forum Call: “Mountains as Sentinels of Change”. The French team (LS, AB, BC, JC, CB, LL, EAH & CLV) was funded by the French National Research Agency (ANR) (ANR-15-MASC-002-01). LS, AB, EAH and CLV were also supported by FEDER InfoBioS (EX011185). LS was partially supported by the German Academic Exchange Service (DAAD) (Short-Term Grant 57440917). The German team (PSY, ST, JM & MTM) was funded by the Deutsche Forschungsgemeinschaft (DFG) (MA 7249/1-1). WC and DWY were supported by the Strategic Priority Research Program of the Chinese Academy of Sciences (XDA20050202), the National Natural Science Foundation of China (41661144002—CLIMTREE grant—, 31670536, 31400470, 31500305), the Key Research Program of Frontier Sciences, CAS (QYZDY-SSW-SMC024), the Bureau of International Cooperation (GJHZ1754), the Ministry of Science and Technology of China (2012FY110800), the State Key Laboratory of Genetic Resources and Evolution (GREKF18-04) at the Kunming Institute of Zoology, the University of East Anglia, and the University of Chinese Academy of Sciences. DF was funded by the Italian CNR—Dipartimento Scienze del Sistema Terra e Tecnologie per l’Ambiente (CNR-DTA) (DTA.AD001.027VB-V).

## Data availability

All scripts and datasets used for analyses are publicly available at the following GitHub repository: https://github.com/Lucasire/Malaise_FR_2017. Raw sequencing data will be available on NCBI upon publication with the following accession number: <u>PRJNA702908</u>.

## Author contributions

This study was conceptualized and designed by CLV, EAH and CB. Forest plot selection and sampling were designed by LL and CB. Sample processing, wet-lab experiments and sequencing were performed by LS, PSY with the help of AB and BC. Bioinformatic demultiplexing was done by LS and PSY with the help of DWY and CW. Ecological and environmental analyses were conducted by LS and WC, with the help of JC, CB, DWY and DF. LS led the writing of the manuscript. All authors contributed substantially to the interpretation and discussion of the results as well as to the revision of the article. All authors approved the submitted version.

