## Supplementary Figures for "Climate-induced forest dieback drives compositional change in insect communities that is concentrated amongst rare species"

Supplementary Fig. 1: **Map of the study area with sampled plot description.**

Map of study area across the three concerned administrative regions of the French Pyrenees. Sampled district are highlighted in pink and blue with plots coloured and shaped according to dieback level and stand type, respectively. Dieback level was assessed using ARCHI method and stand type using PBI measurements.

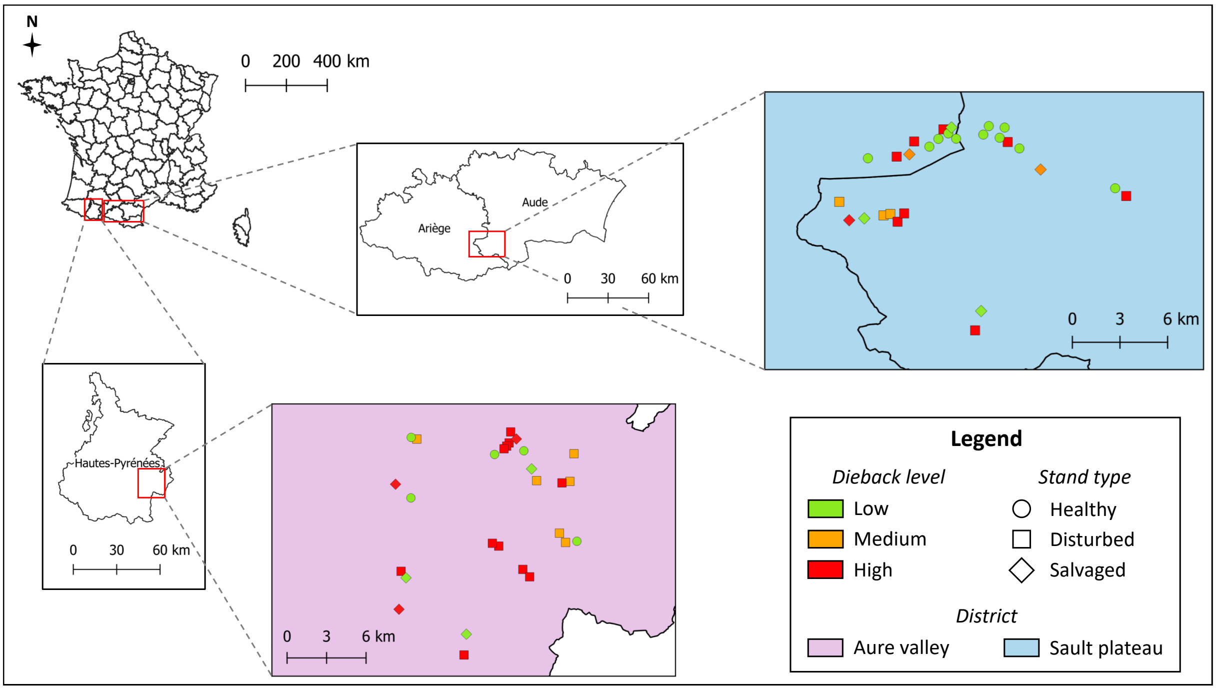

Supplementary Fig. 2: **Observed and estimated insect species richness.**
MOTUs richness observed and estimated, based on Chao2 and Jack1 (first order jackknife) analyses for the 56 plots. While the recovered insect diversity of the present study is of 2972 MOTUs, both estimation methods give around 4000 species trappable using Malaise trap on the four-months sampling period considered.

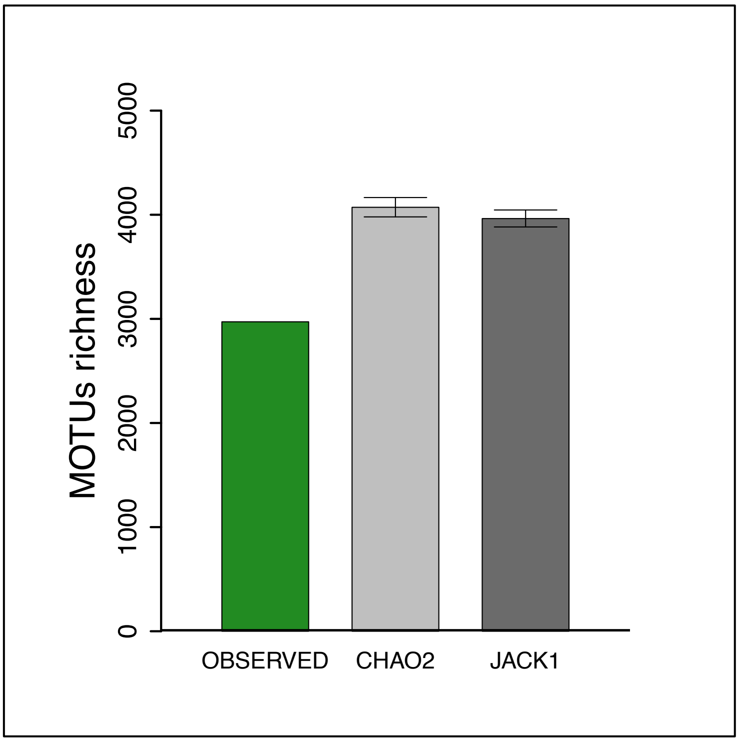

Supplementary Fig. 3: **Relative species prevalence per dieback category and sampled district.**

Heatmaps of MOTU prevalence (proportion of samples in which a species is present) by (**A**) geographical district and (**B**) dieback category. The incidence dataset was bootstrapped 100 times with plot randomization for a total of 5600 virtual samples. Bootstrapped plots were concatenated per (**A**) respective sampled districts or (**B**) dieback categories. For (**A**), prevalence per MOTU is shown per district. Thus, bright yellow (100%) in one district indicates a complete presence of the MOTU in the respective district or in the total dataset. For (**B**), prevalence per MOTU is shown in percentage scaled per dieback category to correspond to 1/3 of the potential 100% maximum presence across all categories. Thus, bright red (33%) in one category means the MOTU is present in 100% of the sampled plots of this respective category. MOTUs can be present at 33% in each category for a total of 100% presence in the complete dataset. Here, we observe that common MOTUs across (**A**) sampled districts or (**B**) dieback categories are the most prevalent and that specific community compositions are mainly shaped by rarely occurring species.

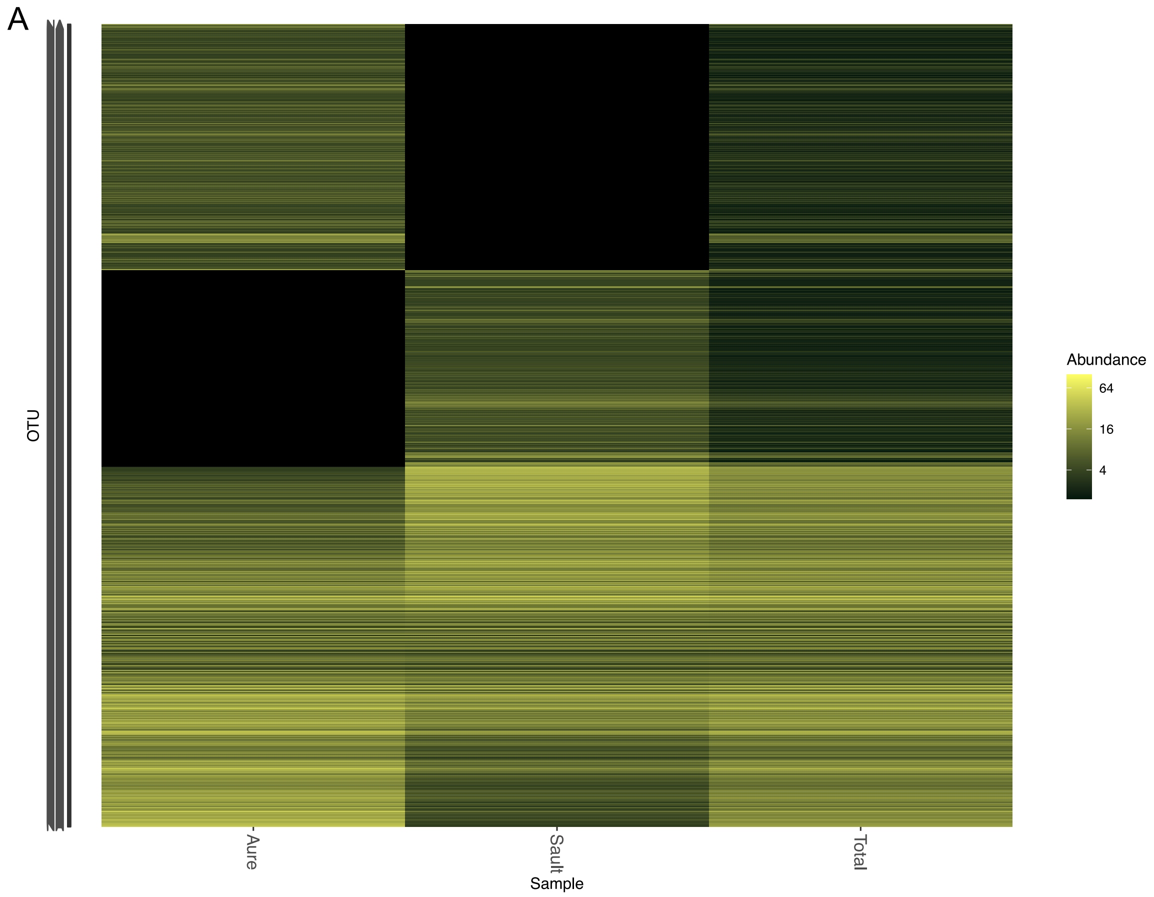

B

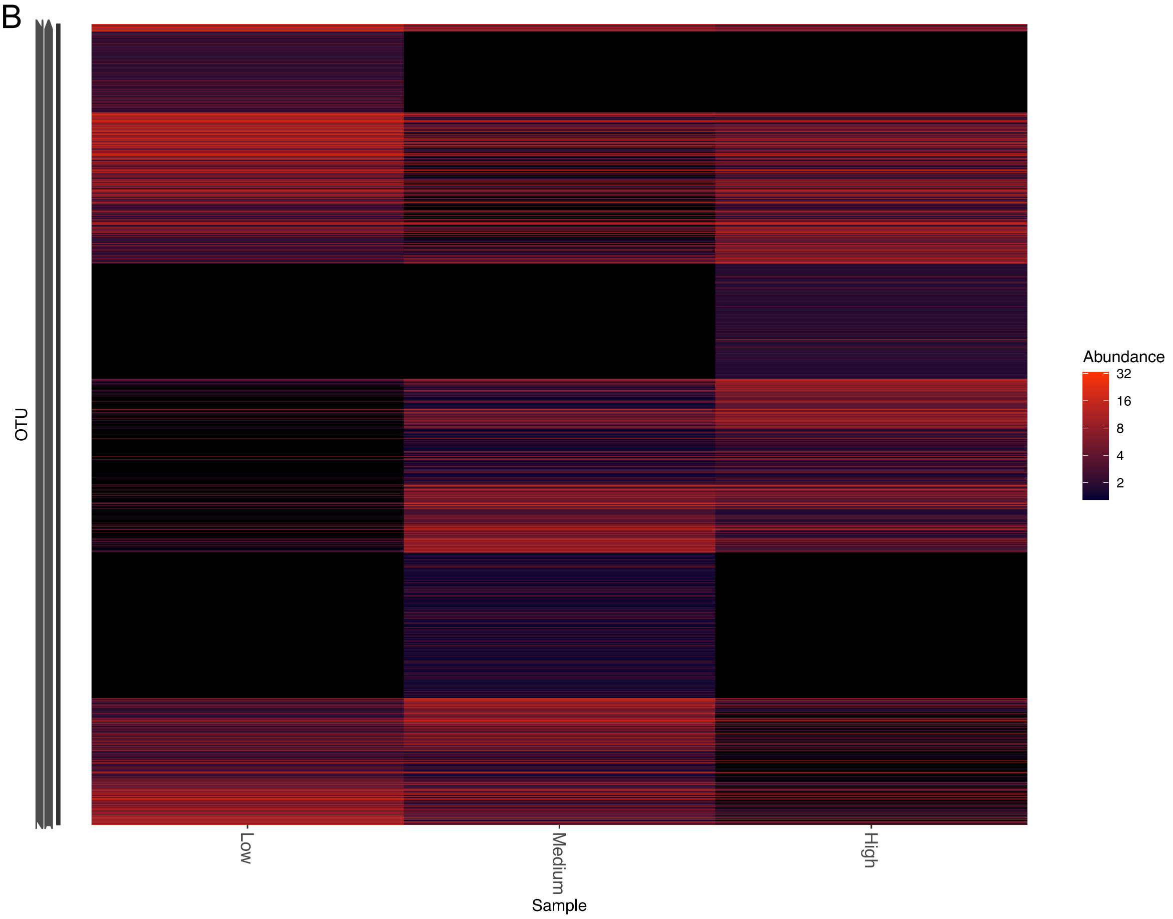

Supplementary Fig. 4: **DNA-based taxonomic identification coverage for the five most represented insect orders.**

Dark green bars show the total number of MOTUs recovered for each of the five most represented insect orders (Diptera, Hymenoptera, Lepidoptera, Coleoptera and Hemiptera). Grey-blue, blue and and light-blue bars represent the number of MOTUs for each order that has been associated with no ambiguity to family, genus and species level, respectively. Taxonomic assignment was performed using BOLD System DNA reference libraries from April 2019.

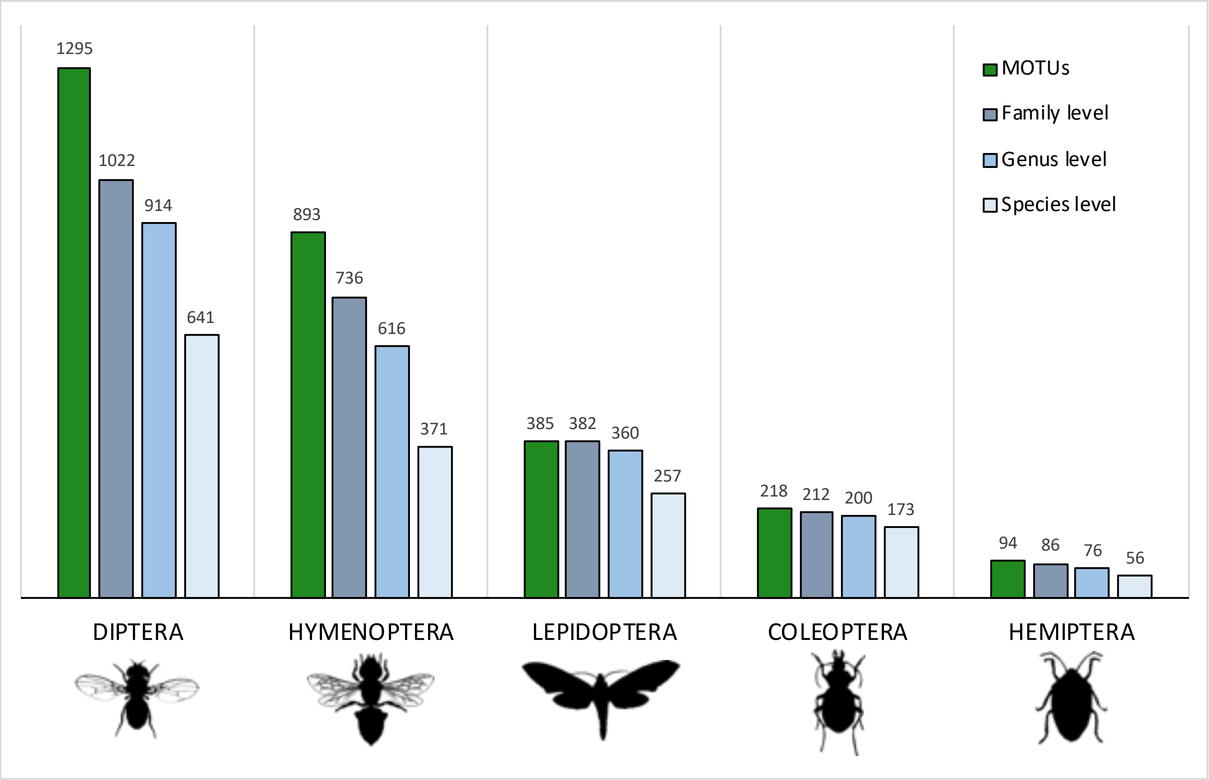

Supplementary Fig. 5: **Zeta-diversity decline and model fitting on the insect fauna for all combinations and assemblages.**

Zeta-diversity analyses per Zeta order (*i.e.* $\zeta_i$) here referring to plots (from $\zeta 2$ to $\zeta 56$) for two computing schemes. Representations consider non-geographically structured scheme that computes all combinations and assemblages (ALL). (**A, B**) Zeta-diversity decline and 0 to 1 scaled ratio of Zeta diversity decline, representing species shared and species retention rate (*i.e.* the retention probability of common species in the community) across $\zeta_i$, respectively. (**C, D**) Zeta-diversity model fitting to exponential and power-law regressions, respectively.

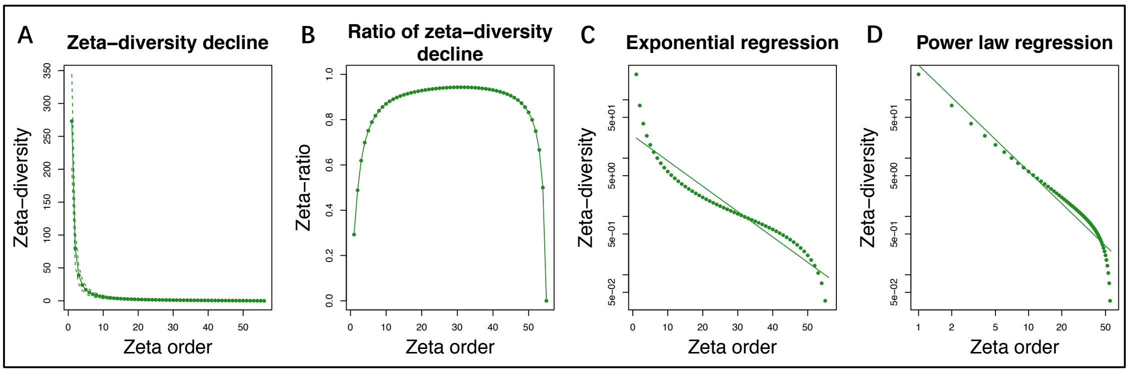

Supplementary Fig. 6: **Effect of dieback on community composition for insect orders and functions.**

Representation of the species retention rate (*i.e.* zeta ratio) per plot (*i.e.* Zeta order) following all plot combinations scheme (ALL) for low, medium and high dieback levels, respectively (**A, B, C**) for the five main insect Orders (Coleoptera, Diptera, Hemiptera, Hymenoptera and Lepidoptera) and (**D, E, F**) for the four main ecological functions recovered from taxonomic assignment (floricolous / non floricolous and parasitoid / non-parasitoid species). Green line with plain dots represents mean species retention rate of the total dataset in each respective dieback category. Increasing curves express that common MOTUs are more likely to be retained in additional samples than rare ones (with presence of common species over all plots if zeta ratio = 1) and decreasing curves indicates species turnover.

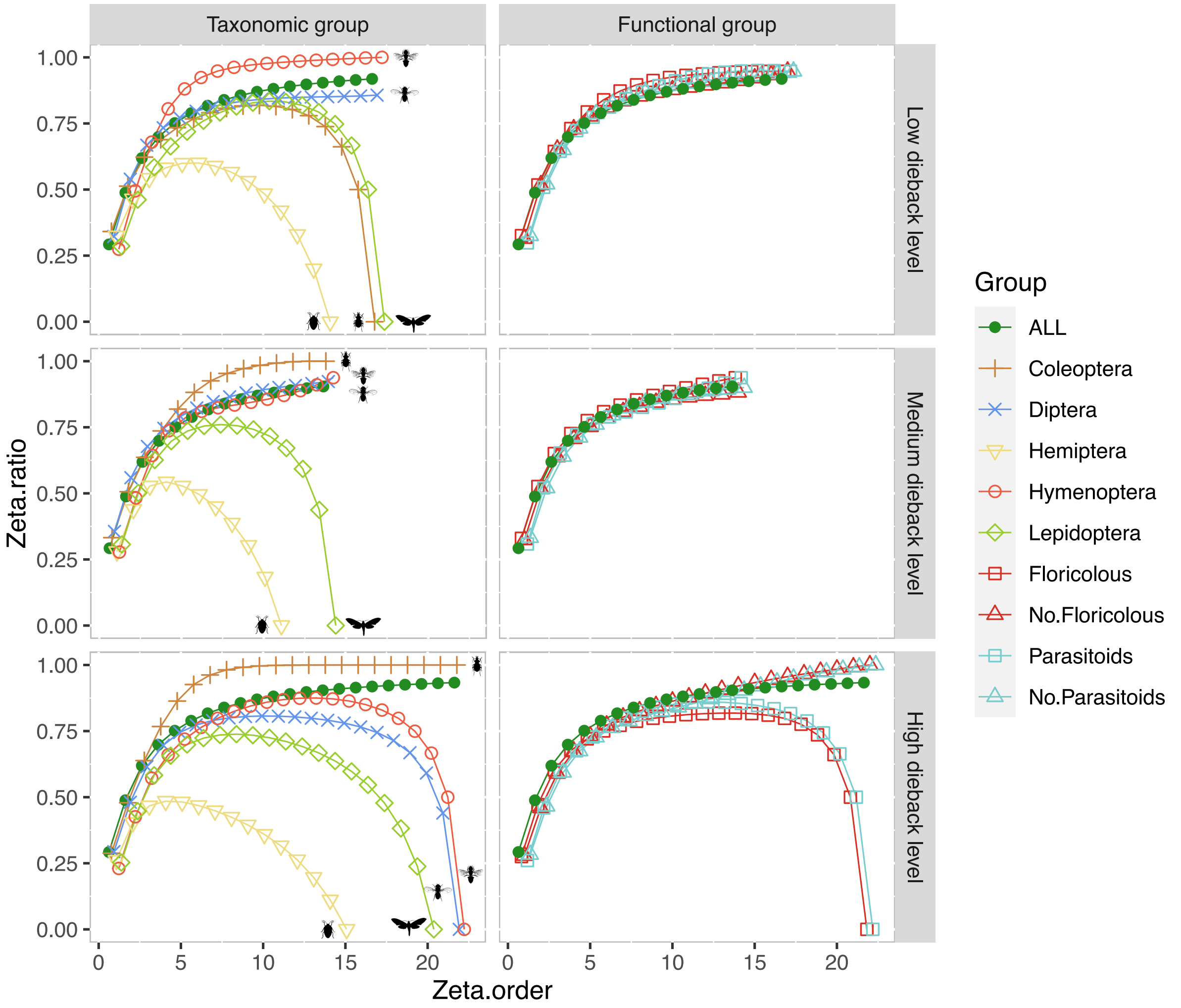

Supplementary Fig. 7: **Effect of stand type on community composition for insect orders and functions.**

Representation of the species retention rate (*i.e.* zeta ratio) per plot (*i.e.* Zeta order) following all plot combinations scheme (ALL) for healthy, disturbed and salvaged stands, respectively (**A, B, C**) for the five main insect Orders (Coleoptera, Diptera, Hemiptera, Hymenoptera and Lepidoptera) and (**D, E, F**) for the four main ecological functions recovered from taxonomic assignment (floricolous / non floricolous and parasitoid / non-parasitoid species). Green line with plain dots represents mean species retention rate of the total dataset in each respective dieback category. Increasing curves express that common MOTUs are more likely to be retained in additional samples than rare ones (with presence of common species over all plots if zeta ratio = 1) and decreasing curves indicates species turnover.

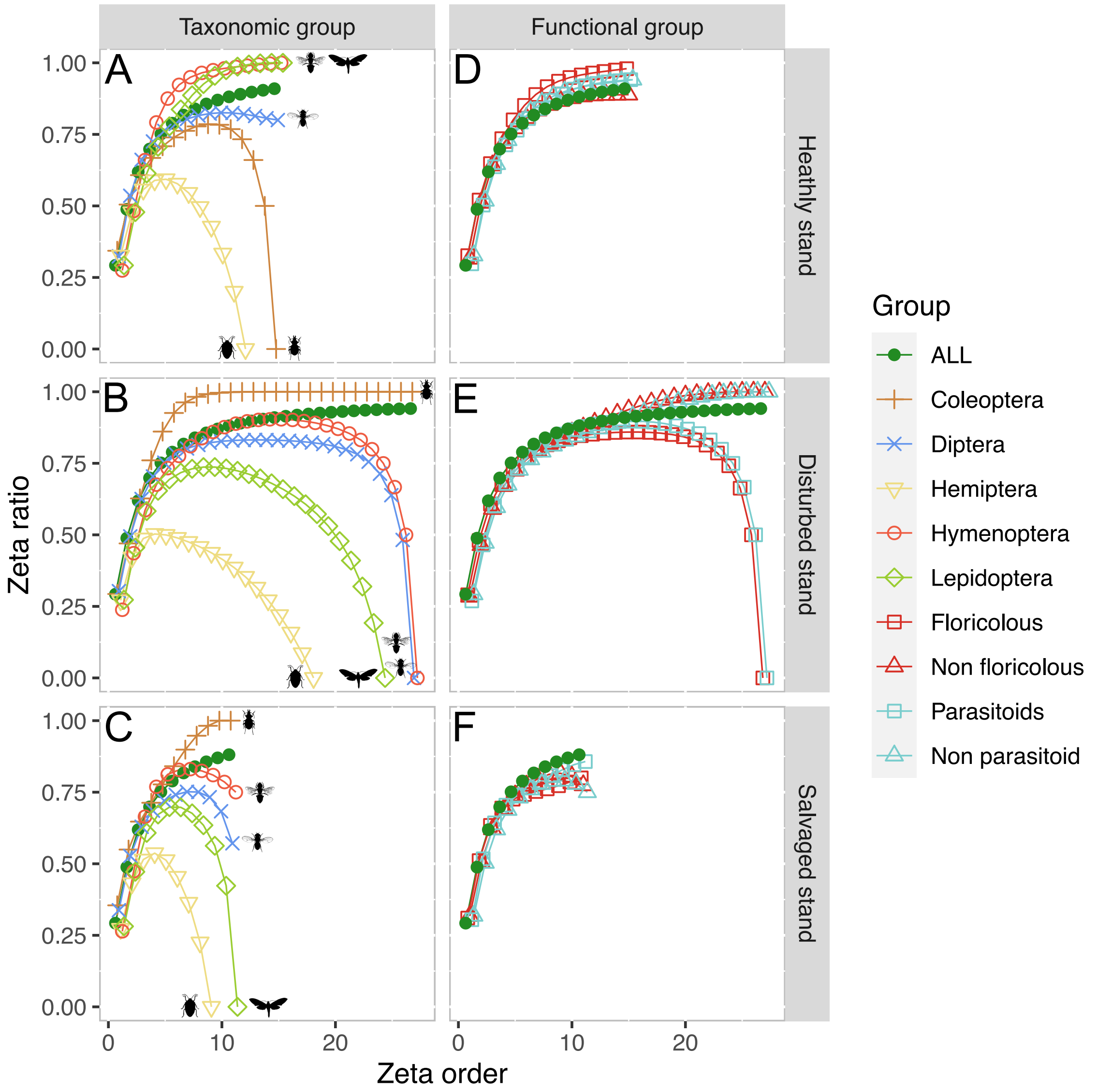
