## Supplementary information - Plot List for "Climate-induced forest dieback drives compositional change in insect communities that is concentrated amongst rare species"

### SI Appendix – Plot list

| Plot | District | Latitude | Longitude | Dieback Level | Stand Type |
|---|---|---|---|---|---|
| ANC4B4 | Aure | 42.874181 | 0.319408 | High | Salvaged |
| ARD3B1 | Aure | 42.919559 | 0.410271 | High | Disturbed |
| ARD3B2 | Aure | 42.921349 | 0.404254 | High | Disturbed |
| ARR3A1 | Aure | 42.99784 | 0.418086 | High | Disturbed |
| ARR4B1 | Aure | 42.993129 | 0.42357 | High | Salvaged |
| BAR1A1 | Aure | 42.929647 | 0.46629 | Medium | Disturbed |
| BAR1A2 | Aure | 42.923449 | 0.472266 | Medium | Disturbed |
| BAR1A3 | Aure | 42.924465 | 0.482811 | Low | Healthy |
| BARE2B1 | Aure | 42.904095 | 0.433298 | High | Disturbed |
| BARE2B2 | Aure | 42.899163 | 0.439834 | High | Disturbed |
| BELGL1A7 | Sault | 42.87165 | 1.96039497 | Low | Healthy |
| BELGL1B5 | Sault | 42.876347 | 1.98102099 | Low | Healthy |
| BELGL1B6 | Sault | 42.876125 | 1.96729898 | Low | Healthy |
| BELGL2B1 | Sault | 42.879281 | 1.97484998 | Low | Healthy |
| BELGL3A8 | Sault | 42.874496 | 1.94864203 | High | Disturbed |
| BELGL3B4 | Sault | 42.881562 | 1.97107804 | High | Disturbed |
| BELGL4B3 | Sault | 42.882689 | 1.97727998 | Medium | Salvaged |
| BEY2B1 | Aure | 42.950613 | 0.327196 | Low | Healthy |
| BEY4B3 | Aure | 42.959642 | 0.312486 | High | Salvaged |
| CAD3B1 | Aure | 42.900214 | 0.320227 | High | Disturbed |
| CAD4B2 | Aure | 42.895868 | 0.324983 | Medium | Salvaged |
| COL1A2 | Sault | 42.883723 | 2.00609998 | Low | Healthy |
| COM1B1 | Sault | 42.877172 | 2.014603 | Low | Healthy |
| COM1B2 | Sault | 42.871258 | 2.02996803 | Low | Healthy |
| COM2B1 | Sault | 42.878847 | 2.00193602 | Low | Healthy |
| COM3B2 | Sault | 42.874816 | 2.02084098 | High | Disturbed |
| COM4B1.1 | Sault | 42.859428 | 2.04649899 | Low | Salvaged |
| COMU1A1 | Sault | 42.839738 | 1.89150402 | Medium | Disturbed |
| COMU2A1 | Sault | 42.832367 | 1.925698 | Medium | Disturbed |
| COMU2A2 | Sault | 42.833225 | 1.93099503 | Medium | Disturbed |
| COMU3A1 | Sault | 42.828883 | 1.936599 | High | Disturbed |
| COMU3B1 | Sault | 42.833637 | 1.94162797 | High | Disturbed |
| COMU4B1.1 | Sault | 42.829366 | 1.89930397 | High | Salvaged |
| COMU4B1.3 | Sault | 42.830545 | 1.91089899 | Medium | Salvaged |
| FOU1A1 | Sault | 42.864636 | 1.91310796 | Low | Healthy |
| FOU2A1 | Sault | 42.865748 | 1.93526996 | High | Disturbed |
| FOU4B2.1 | Sault | 42.867145 | 1.94505097 | Low | Salvaged |
| GOU3B1 | Aure | 42.844392 | 0.381135 | High | Disturbed |

| | | | | | |
|---|---|---|---|---|---|
| GOU4B1 | Aure | 42.858618 | 0.382687 | Medium | Salvaged |
| HEC3A1 | Aure | 42.990304 | 0.416893 | High | Disturbed |
| HECGF1A3 | Aure | 42.991953 | 0.325686 | Low | Healthy |
| HECGF2A2 | Aure | 42.990933 | 0.330944 | Medium | Disturbed |
| ILH2A1 | Aure | 42.988058 | 0.414755 | High | Disturbed |
| MER2B1 | Sault | 42.767846 | 1.99743301 | High | Disturbed |
| MER4B1.2 | Sault | 42.778858 | 2.00187701 | Medium | Salvaged |
| NISE1A1 | Aure | 42.984276 | 0.477635 | Medium | Disturbed |
| NISE1A2 | Aure | 42.965282 | 0.474635 | Medium | Disturbed |
| NISE1A3 | Aure | 42.964101 | 0.467242 | High | Disturbed |
| NISW1A1 | Aure | 42.985233 | 0.430842 | Low | Healthy |
| NISW1A2 | Aure | 42.965114 | 0.443573 | Medium | Disturbed |
| NISW2A1 | Aure | 42.844962 | 2.11278803 | High | Disturbed |
| PEY1B1 | Sault | 42.982117 | 0.403709 | Low | Healthy |
| PEYR3A1 | Sault | 42.986194 | 0.412343 | High | Disturbed |
| PUI1A1 | Sault | 42.882982 | 2.01833303 | Low | Healthy |
| SAR1A3 | Aure | 42.849289 | 2.10417101 | Low | Healthy |
| SAR4B1 | Aure | 42.973043 | 0.438573 | Medium | Salvaged |
