## Supplementary information - Primer List for "Climate-induced forest dieback drives compositional change in insect communities that is concentrated amongst rare species"

### SI Appendix – Primer list

| **Forward primers** | | **Reverse primers** | |
|---|---|---|---|
| ***Name*** | ***Tagged forward primer 5′-3′*** | ***Name*** | ***Tagged reverse primer 5′-3′*** |
| F1_Leray | ATACAGTCGGWACWGGWTGAACWGTWTAYCCYCC | R1_Leray | ATACAGTCTAIACYTCIGGRTGICCRAARAAYCA |
| F2_Leray | AGCCTTAAGGWACWGGWTGAACWGTWTAYCCYCC | R2_Leray | AGCCTTAATAIACYTCIGGRTGICCRAARAAYCA |
| F3_Leray | ATTAGCCTGGWACWGGWTGAACWGTWTAYCCYCC | R3_Leray | ATTAGCCTTAIACYTCIGGRTGICCRAARAAYCA |
| F4_Leray | ATTAGGAAGGWACWGGWTGAACWGTWTAYCCYCC | R4_Leray | ATTAGGAATAIACYTCIGGRTGICCRAARAAYCA |
| F5_Leray | ATCTACGAGGWACWGGWTGAACWGTWTAYCCYCC | R5_Leray | ATCTACGATAIACYTCIGGRTGICCRAARAAYCA |
| F6_Leray | ACGTCTTAGGWACWGGWTGAACWGTWTAYCCYCC | R6_Leray | ACGTCTTATAIACYTCIGGRTGICCRAARAAYCA |
| F7_Leray | ATTAGTTCGGWACWGGWTGAACWGTWTAYCCYCC | R7_Leray | ATTAGTTCTAIACYTCIGGRTGICCRAARAAYCA |
| F8_Leray | ATTATAATGGWACWGGWTGAACWGTWTAYCCYCC | R8_Leray | ATTATAATTAIACYTCIGGRTGICCRAARAAYCA |
| F9_Leray | AGGTTCTAGGWACWGGWTGAACWGTWTAYCCYCC | R9_Leray | AGGTTCTATAIACYTCIGGRTGICCRAARAAYCA |
| F10_Leray | ACTTGATCGGWACWGGWTGAACWGTWTAYCCYCC | R10_Leray | ACTTGATCTAIACYTCIGGRTGICCRAARAAYCA |
| F11_Leray | ATCTCATAGGWACWGGWTGAACWGTWTAYCCYCC | R11_Leray | ATCTCATATAIACYTCIGGRTGICCRAARAAYCA |
| F12_Leray | ACATTCTCGGWACWGGWTGAACWGTWTAYCCYCC | R12_Leray | ACATTCTCTAIACYTCIGGRTGICCRAARAAYCA |
| F13_Leray | ATCTCTATGGWACWGGWTGAACWGTWTAYCCYCC | R13_Leray | ATCTCTATTAIACYTCIGGRTGICCRAARAAYCA |
| F14_Leray | ACATTGGAGGWACWGGWTGAACWGTWTAYCCYCC | R14_Leray | ACATTGGATAIACYTCIGGRTGICCRAARAAYCA |
| F15_Leray | AGAACAGTGGWACWGGWTGAACWGTWTAYCCYCC | R15_Leray | AGAACAGTTAIACYTCIGGRTGICCRAARAAYCA |
| F16_Leray | ATAGAGCAGGWACWGGWTGAACWGTWTAYCCYCC | R16_Leray | ATAGAGCATAIACYTCIGGRTGICCRAARAAYCA |
| F17_Leray | ATCTGAGTGGWACWGGWTGAACWGTWTAYCCYCC | R17_Leray | ATCTGAGTTAIACYTCIGGRTGICCRAARAAYCA |
| F18_Leray | ATTCAAGCGGWACWGGWTGAACWGTWTAYCCYCC | R18_Leray | ATTCAAGCTAIACYTCIGGRTGICCRAARAAYCA |
| F19_Leray | ATTCACCAGGWACWGGWTGAACWGTWTAYCCYCC | R19_Leray | ATTCACCATAIACYTCIGGRTGICCRAARAAYCA |
| F20_Leray | ATTCAGATGGWACWGGWTGAACWGTWTAYCCYCC | R20_Leray | ATTCAGATTAIACYTCIGGRTGICCRAARAAYCA |
| F21_Leray | ATAGCATCGGWACWGGWTGAACWGTWTAYCCYCC | R21_Leray | ATAGCATCTAIACYTCIGGRTGICCRAARAAYCA |

| | | | |
|---|---|---|---|
| F22_Leray | ATGTTGGTGGWACWGGWTGAACWGTWTAYCCYCC | R22_Leray | ATGTTGGTTAIACYTCIGGRTGICCRAARAAYCA |
| F23_Leray | ATCCAACTGGWACWGGWTGAACWGTWTAYCCYCC | R23_Leray | ATCCAACTTAIACYTCIGGRTGICCRAARAAYCA |
| F24_Leray | ATAATGCTGGWACWGGWTGAACWGTWTAYCCYCC | R24_Leray | ATAATGCTTAIACYTCIGGRTGICCRAARAAYCA |
| F25_Leray | ATAGTCGTGGWACWGGWTGAACWGTWTAYCCYCC | R25_Leray | ATAGTCGTTAIACYTCIGGRTGICCRAARAAYCA |
| F26_Leray | CAGCGAATAGGWACWGGWTGAACWGTWTAYCCYCC | R26_Leray | CAGCGAATATAIACYTCIGGRTGICCRAARAAYCA |
| F27_Leray | CACTCCATAGGWACWGGWTGAACWGTWTAYCCYCC | R27_Leray | CACTCCATATAIACYTCIGGRTGICCRAARAAYCA |
| F28_Leray | CAGCTACATGGWACWGGWTGAACWGTWTAYCCYCC | R28_Leray | CAGCTACATTAIACYTCIGGRTGICCRAARAAYCA |
| F29_Leray | CAGCTATCAGGWACWGGWTGAACWGTWTAYCCYCC | R29_Leray | CAGCTATCATAIACYTCIGGRTGICCRAARAAYCA |
| F30_Leray | CACTGAACAGGWACWGGWTGAACWGTWTAYCCYCC | R30_Leray | CACTGAACATAIACYTCIGGRTGICCRAARAAYCA |
| F31_Leray | CATGTGTCTGGWACWGGWTGAACWGTWTAYCCYCC | R31_Leray | CATGTGTCTTAIACYTCIGGRTGICCRAARAAYCA |
| F32_Leray | CACTGACATGGWACWGGWTGAACWGTWTAYCCYCC | R32_Leray | CACTGACATTAIACYTCIGGRTGICCRAARAAYCA |
| F33_Leray | CATCCGTTAGGWACWGGWTGAACWGTWTAYCCYCC | R33_Leray | CATCCGTTATAIACYTCIGGRTGICCRAARAAYCA |
| F34_Leray | CATCATGTAGGWACWGGWTGAACWGTWTAYCCYCC | R34_Leray | CATCATGTATAIACYTCIGGRTGICCRAARAAYCA |
| F35_Leray | CATCCTTGTGGWACWGGWTGAACWGTWTAYCCYCC | R35_Leray | CATCCTTGTTAIACYTCIGGRTGICCRAARAAYCA |
| F36_Leray | CATTAATGTGGWACWGGWTGAACWGTWTAYCCYCC | R36_Leray | CATTAATGTTAIACYTCIGGRTGICCRAARAAYCA |
| F37_Leray | CATTACACCGGWACWGGWTGAACWGTWTAYCCYCC | R37_Leray | CATTACACCTAIACYTCIGGRTGICCRAARAAYCA |
| F38_Leray | CATGTTACAGGWACWGGWTGAACWGTWTAYCCYCC | R38_Leray | CATGTTACATAIACYTCIGGRTGICCRAARAAYCA |
| F39_Leray | CATTACCGAGGWACWGGWTGAACWGTWTAYCCYCC | R39_Leray | CATTACCGATAIACYTCIGGRTGICCRAARAAYCA |
| F40_Leray | CACTGTTGTGGWACWGGWTGAACWGTWTAYCCYCC | R40_Leray | CACTGTTGTTAIACYTCIGGRTGICCRAARAAYCA |
| F41_Leray | CGTAAGTGAGGWACWGGWTGAACWGTWTAYCCYCC | R41_Leray | CGTAAGTGATAIACYTCIGGRTGICCRAARAAYCA |
| F42_Leray | CGTACCAGAGGWACWGGWTGAACWGTWTAYCCYCC | R42_Leray | CGTACCAGATAIACYTCIGGRTGICCRAARAAYCA |
| F43_Leray | CGCTCGTATGGWACWGGWTGAACWGTWTAYCCYCC | R43_Leray | CGCTCGTATTAIACYTCIGGRTGICCRAARAAYCA |
| F44_Leray | CGCTCTAACGGWACWGGWTGAACWGTWTAYCCYCC | R44_Leray | CGCTCTAACTAIACYTCIGGRTGICCRAARAAYCA |
| F45_Leray | CGTCATTCCGGWACWGGWTGAACWGTWTAYCCYCC | R45_Leray | CGTCATTCCTAIACYTCIGGRTGICCRAARAAYCA |
| F46_Leray | CGTAATAGCGGWACWGGWTGAACWGTWTAYCCYCC | R46_Leray | CGTAATAGCTAIACYTCIGGRTGICCRAARAAYCA |
| F47_Leray | CGTGTTCACGGWACWGGWTGAACWGTWTAYCCYCC | R47_Leray | CGTGTTCACTAIACYTCIGGRTGICCRAARAAYCA |
| F48_Leray | CGTAATCAAGGWACWGGWTGAACWGTWTAYCCYCC | R48_Leray | CGTAATCAATAIACYTCIGGRTGICCRAARAAYCA |
| F49_Leray | CGTCCATACGGWACWGGWTGAACWGTWTAYCCYCC | R49_Leray | CGTCCATACTAIACYTCIGGRTGICCRAARAAYCA |

| | | | |
|---|---|---|---|
| F50_Leray | CGTTAACACGGWACWGGWTGAACWGTWTAYCCYCC | R50_Leray | CGTTAACACTAIACYTCIGGRTGICCRAARAAYCA |
| F51_Leray | GCACGTTACGGWACWGGWTGAACWGTWTAYCCYCC | R51_Leray | GCACGTTACTAIACYTCIGGRTGICCRAARAAYCA |
| F52_Leray | GCAACAGGTGGWACWGGWTGAACWGTWTAYCCYCC | R52_Leray | GCAACAGGTTAIACYTCIGGRTGICCRAARAAYCA |
| F53_Leray | GCACTCACTGGWACWGGWTGAACWGTWTAYCCYCC | R53_Leray | GCACTCACTTAIACYTCIGGRTGICCRAARAAYCA |
| F54_Leray | GCAACTCCTGGWACWGGWTGAACWGTWTAYCCYCC | R54_Leray | GCAACTCCTTAIACYTCIGGRTGICCRAARAAYCA |
| F55_Leray | GCAAGAAGCGGWACWGGWTGAACWGTWTAYCCYCC | R55_Leray | GCAAGAAGCTAIACYTCIGGRTGICCRAARAAYCA |
| F56_Leray | GCATCATCTGGWACWGGWTGAACWGTWTAYCCYCC | R56_Leray | GCATCATCTTAIACYTCIGGRTGICCRAARAAYCA |
| F57_Leray | GCAAGATATGGWACWGGWTGAACWGTWTAYCCYCC | R57_Leray | GCAAGATATTAIACYTCIGGRTGICCRAARAAYCA |
| F58_Leray | GCATCCTACGGWACWGGWTGAACWGTWTAYCCYCC | R58_Leray | GCATCCTACTAIACYTCIGGRTGICCRAARAAYCA |
| F59_Leray | GCATGTGCTGGWACWGGWTGAACWGTWTAYCCYCC | R59_Leray | GCATGTGCTTAIACYTCIGGRTGICCRAARAAYCA |
| F60_Leray | GCAGAGGTAGGWACWGGWTGAACWGTWTAYCCYCC | R60_Leray | GCAGAGGTATAIACYTCIGGRTGICCRAARAAYCA |
| F61_Leray | GTAACAACAGGWACWGGWTGAACWGTWTAYCCYCC | R61_Leray | GTAACAACATAIACYTCIGGRTGICCRAARAAYCA |
| F62_Leray | GTAACACACGGWACWGGWTGAACWGTWTAYCCYCC | R62_Leray | GTAACACACTAIACYTCIGGRTGICCRAARAAYCA |
| F63_Leray | GTACTAATCGGWACWGGWTGAACWGTWTAYCCYCC | R63_Leray | GTACTAATCTAIACYTCIGGRTGICCRAARAAYCA |
| F64_Leray | GTAACCGTAGGWACWGGWTGAACWGTWTAYCCYCC | R64_Leray | GTAACCGTATAIACYTCIGGRTGICCRAARAAYCA |
| F65_Leray | GTAACGATCGGWACWGGWTGAACWGTWTAYCCYCC | R65_Leray | GTAACGATCTAIACYTCIGGRTGICCRAARAAYCA |
| F66_Leray | GTAACGTAAGGWACWGGWTGAACWGTWTAYCCYCC | R66_Leray | GTAACGTAATAIACYTCIGGRTGICCRAARAAYCA |
| F67_Leray | GTGCTTAGAGGWACWGGWTGAACWGTWTAYCCYCC | R67_Leray | GTGCTTAGATAIACYTCIGGRTGICCRAARAAYCA |
| F68_Leray | GTCTGATTCGGWACWGGWTGAACWGTWTAYCCYCC | R68_Leray | GTCTGATTCTAIACYTCIGGRTGICCRAARAAYCA |
| F69_Leray | GTCTGCTAAGGWACWGGWTGAACWGTWTAYCCYCC | R69_Leray | GTCTGCTAATAIACYTCIGGRTGICCRAARAAYCA |
| F70_Leray | GTACTGCTAGGWACWGGWTGAACWGTWTAYCCYCC | R70_Leray | GTACTGCTATAIACYTCIGGRTGICCRAARAAYCA |
| F71_Leray | GTCACTTCAGGWACWGGWTGAACWGTWTAYCCYCC | R71_Leray | GTCACTTCATAIACYTCIGGRTGICCRAARAAYCA |
| F72_Leray | GTAACCAATGGWACWGGWTGAACWGTWTAYCCYCC | R72_Leray | GTAACCAATTAIACYTCIGGRTGICCRAARAAYCA |
| F73_Leray | GTCTTAAGTGGWACWGGWTGAACWGTWTAYCCYCC | R73_Leray | GTCTTAAGTTAIACYTCIGGRTGICCRAARAAYCA |
| F74_Leray | GTCTCATGAGGWACWGGWTGAACWGTWTAYCCYCC | R74_Leray | GTCTCATGATAIACYTCIGGRTGICCRAARAAYCA |
| F75_Leray | GTCTTAGAAGGWACWGGWTGAACWGTWTAYCCYCC | R75_Leray | GTCTTAGAATAIACYTCIGGRTGICCRAARAAYCA |
| F76_Leray | TGGAATATGGWACWGGWTGAACWGTWTAYCCYCC | R76_Leray | TGGAATATTAIACYTCIGGRTGICCRAARAAYCA |
| F77_Leray | TACTTGCAGGWACWGGWTGAACWGTWTAYCCYCC | R77_Leray | TACTTGCATAIACYTCIGGRTGICCRAARAAYCA |

| | |
|---|---|
| F78_Leray | TGAAGCATGGWACWGGWTGAACWGTWTAYCCYCC |
| F79_Leray | TCACTAGTGGWACWGGWTGAACWGTWTAYCCYCC |
| F80_Leray | TAAGCTCAGGWACWGGWTGAACWGTWTAYCCYCC |
| F81_Leray | TAGAACGTGGWACWGGWTGAACWGTWTAYCCYCC |
| F82_Leray | TAGAAGACGGWACWGGWTGAACWGTWTAYCCYCC |
| F83_Leray | TAAGGACTGGWACWGGWTGAACWGTWTAYCCYCC |
| F84_Leray | TAGAATCAGGWACWGGWTGAACWGTWTAYCCYCC |
| F85_Leray | TAAGTGAAGGWACWGGWTGAACWGTWTAYCCYCC |
| F86_Leray | TAGTCGATGGWACWGGWTGAACWGTWTAYCCYCC |
| F87_Leray | TGAACTAAGGWACWGGWTGAACWGTWTAYCCYCC |
| F88_Leray | TAGATACCGGWACWGGWTGAACWGTWTAYCCYCC |
| F89_Leray | TAATAGCCGGWACWGGWTGAACWGTWTAYCCYCC |
| F90_Leray | TAATATGAGGWACWGGWTGAACWGTWTAYCCYCC |
| F91_Leray | TATGCAGTGGWACWGGWTGAACWGTWTAYCCYCC |
| F92_Leray | TAATCCAAGGWACWGGWTGAACWGTWTAYCCYCC |
| F93_Leray | TGAATATAGGWACWGGWTGAACWGTWTAYCCYCC |
| F94_Leray | TATATCCAGGWACWGGWTGAACWGTWTAYCCYCC |
| F95_Leray | TGAATGACGGWACWGGWTGAACWGTWTAYCCYCC |
| F96_Leray | TAGCATTCGGWACWGGWTGAACWGTWTAYCCYCC |
| F97_Leray | TAATCTTCGGWACWGGWTGAACWGTWTAYCCYCC |
| F98_Leray | TGACAGAAGGWACWGGWTGAACWGTWTAYCCYCC |
| F99_Leray | TCCTGTAAGGWACWGGWTGAACWGTWTAYCCYCC |
| F100_Leray | TAGCTCAAGGWACWGGWTGAACWGTWTAYCCYCC |

| | |
|---|---|
| R78_Leray | TGAAGCATTAIACYTCIGGRTGICCRAARAAYCA |
| R79_Leray | TCACTAGTTAIACYTCIGGRTGICCRAARAAYCA |
| R80_Leray | TAAGCTCATAIACYTCIGGRTGICCRAARAAYCA |
| R81_Leray | TAGAACGTTAIACYTCIGGRTGICCRAARAAYCA |
| R82_Leray | TAGAAGACTAIACYTCIGGRTGICCRAARAAYCA |
| R83_Leray | TAAGGACTTAIACYTCIGGRTGICCRAARAAYCA |
| R84_Leray | TAGAATCATAIACYTCIGGRTGICCRAARAAYCA |
| R85_Leray | TAAGTGAATAIACYTCIGGRTGICCRAARAAYCA |
| R86_Leray | TAGTCGATTAIACYTCIGGRTGICCRAARAAYCA |
| R87_Leray | TGAACTAATAIACYTCIGGRTGICCRAARAAYCA |
| R88_Leray | TAGATACCTAIACYTCIGGRTGICCRAARAAYCA |
| R89_Leray | TAATAGCCTAIACYTCIGGRTGICCRAARAAYCA |
| R90_Leray | TAATATGATAIACYTCIGGRTGICCRAARAAYCA |
| R91_Leray | TATGCAGTTAIACYTCIGGRTGICCRAARAAYCA |
| R92_Leray | TAATCCAATAIACYTCIGGRTGICCRAARAAYCA |
| R93_Leray | TGAATATATAIACYTCIGGRTGICCRAARAAYCA |
| R94_Leray | TATATCCATAIACYTCIGGRTGICCRAARAAYCA |
| R95_Leray | TGAATGACTAIACYTCIGGRTGICCRAARAAYCA |
| R96_Leray | TAGCATTCTAIACYTCIGGRTGICCRAARAAYCA |
| R97_Leray | TAATCTTCTAIACYTCIGGRTGICCRAARAAYCA |
| R98_Leray | TGACAGAATAIACYTCIGGRTGICCRAARAAYCA |
| R99_Leray | TCCTGTAATAIACYTCIGGRTGICCRAARAAYCA |
| R100_Leray | TAGCTCAATAIACYTCIGGRTGICCRAARAAYCA |
