## Supplementary Table II for "Climate-induced forest dieback drives compositional change in insect communities that is concentrated amongst rare species"

Supplementary Table II: **Impact of forest dieback and salvage logging on insect communities.**

Generalized linear models from mvabund object for community composition changes of different study groups (i.e. total insects, five most represented insect orders, and four functional group) compared across respective environmental conditions of diebacks (i.e. low, medium and high forest dieback levels) and stand types (i.e. healthy, disturbed and salvaged logged). Functional groups (i.e. Parasitoids/non-parasitoids, floricolous/non-floricolous insects) were assigned using each MOTU's taxonomic family. Reported values are the two extremes given from 10 consecutive analyses. Post-Hoc Holm correction was applied on the complete 10 trials range. Significance is given by "*" with relatives values in bold while "N.S." stands for non-significant. We highlight that all studied groups but Coleoptera had community composition changes driven forest dieback. Significant compositional changes also arose in each cases between study districts.

| Studied group | Variable effect | Effect' range of p-values (95% confidence) | Post-Hoc Holm correction | Significance |
|---|---|---|---|---|
| **Total insects** | | | | |
| | Forest dieback | 0.007 to 0.001 | **0.024 to 0.018** | * to ** |
| | Stand type | 0.062 to 0.049 | 0.49 | *N.S.* |
| | District | 0.001 | **0.01** | ** |
| **Coleoptera** | | | | |
| | Forest dieback | 0.043 to 0.024 | 0.27 | *N.S.* |
| | Stand type | 0.210 to 0.163 | 1 | *N.S.* |
| | District | 0.001 | **0.01** | ** |
| **Diptera** | | | | |
| | Forest dieback | 0.005 to 0.002 | **0.028 to to 0.02** | * |
| | Stand type | 0.149 to 0.109 | 1 | *N.S.* |
| | District | 0.001 | **0.01** | ** |
| **Hemiptera** | | | | |
| | Forest dieback | 0.008 to 0.002 | **0.036 to 0.02** | * |
| | Stand type | 0.205 to 0.176 | 1 | *N.S.* |
| | District | 0.001 | **0.01** | ** |
| **Hymenoptera** | | | | |
| | Forest dieback | 0.007 to 0.001 | **0.03 to 0.01** | * to ** |
| | Stand type | 0.042 to 0.027 | 0.28 | *N.S.* |
| | District | 0.001 | **0.01** | ** |
| **Lepidoptera** | | | | |
| | Forest dieback | 0.011 to 0.003 | 0.064 to **0.03** | *N.S.* to * |
| | Stand type | 0.025 to 0.009 | 0.144 to 0.09 | *N.S.* |
| | District | 0.001 | **0.01** | ** |
| **Floricolous** | | | | |
| | Forest dieback | 0.007 to 0.003 | **0.03** | * |
| | Stand type | 0.056 to 0.038 | 0.38 | *N.S.* |
| | District | 0.001 | **0.01** | ** |
| **Non-floricolous** | | | | |
| | Forest dieback | 0.004 to 0.001 | **0.016** to **0.01** | ** |
| | Stand type | 0.08 to 0.06 | 0.6 | *N.S.* |
| | District | 0.001 | **0.01** | ** |
| **Parasitoids** | | | | |
| | Forest dieback | 0.008 to 0.001 | **0.024** to **0.01** | * to ** |
| | Stand type | 0.033 to 0.016 | 0.26 to 0.16 | *N.S.* |
| | District | 0.001 | **0.01** | ** |
| **Non-parasitoids** | | | | |
| | Forest dieback | 0.006 to 0.002 | **0.024** to **0.02** | * |
| | Stand type | 0.091 to 0.063 | 0.63 | *N.S.* |
| | District | 0.001 | **0.01** | ** |
