## Supplementary Table III for "Climate-induced forest dieback drives compositional change in insect communities that is concentrated amongst rare species"

Supplementary Table III: **Species associated with a specific dieback level**

Table of IndVal analysis to determine OTUs significantly associated to each dieback level (*i.e.* low, medium and high). IndVal analyses were performed 10 times and only OTUs with 10 significant associations for each dieback level were retained. Post-hoc Holm correction was then applied on the 10 p-values range reported for each OTU. Significant p-values after Holm correction are highlighted in bold. *N.S.* stands for non significant.

| Taxonomy | OTUs | Uncorrected | | | | | | | | | | Holm correction | | | | | | | | | | Significance | Dieback level |
|---|---|---|---|---|---|---|---|---|---|---|---|---|---|---|---|---|---|---|---|---|---|---|---|
| | | RUN 1 | RUN 2 | RUN 3 | RUN 4 | RUN 5 | RUN 6 | RUN 7 | RUN 8 | RUN 9 | RUN 10 | RUN 1 | RUN 2 | RUN 3 | RUN 4 | RUN 5 | RUN 6 | RUN 7 | RUN 8 | RUN 9 | RUN 10 | | |
| Entiminae sp. | OTU1110 | 0.025 | 0.016 | 0.019 | 0.024 | 0.018 | 0.019 | 0.027 | 0.021 | 0.025 | 0.026 | 0.162 | 0.16 | 0.162 | 0.162 | 0.162 | 0.162 | 0.162 | 0.162 | 0.162 | 0.162 | N.S. | LOW |
| Peyerimhoffina gracilis | OTU1115 | 0.005 | 0.005 | 0.003 | 0.005 | 0.003 | 0.01 | 0.002 | 0.007 | 0.003 | 0.002 | **0.025** | **0.025** | **0.024** | **0.025** | **0.024** | **0.025** | **0.02** | **0.025** | **0.024** | **0.02** | * to ** | |
| Cecidomyiidae sp. | OTU1127 | 0.002 | 0.001 | 0.001 | 0.002 | 0.003 | 0.004 | 0.002 | 0.005 | 0.005 | 0.003 | **0.016** | **0.01** | **0.01** | **0.016** | **0.016** | **0.016** | **0.016** | **0.016** | **0.016** | **0.016** | ** | |
| Diptera sp. | OTU1155 | 0.005 | 0.004 | 0.005 | 0.005 | 0.008 | 0.004 | 0.005 | 0.009 | 0.009 | 0.01 | **0.04** | **0.04** | **0.04** | **0.04** | **0.04** | **0.04** | **0.04** | **0.04** | **0.04** | **0.04** | * | |
| Diptera sp. | OTU1168 | 0.012 | 0.012 | 0.014 | 0.016 | 0.017 | 0.012 | 0.015 | 0.009 | 0.008 | 0.015 | 0.096 | 0.096 | 0.096 | 0.096 | 0.096 | 0.096 | 0.096 | 0.081 | 0.08 | 0.096 | N.S. | |
| Enclisis ornaticeps | OTU1169 | 0.022 | 0.025 | 0.011 | 0.021 | 0.019 | 0.019 | 0.012 | 0.018 | 0.028 | 0.025 | 0.144 | 0.144 | 0.11 | 0.144 | 0.144 | 0.144 | 0.11 | 0.144 | 0.144 | 0.144 | N.S. | |
| Hyperlasion wasmanni | OTU120 | 0.003 | 0.002 | 0.002 | 0.005 | 0.005 | 0.003 | 0.002 | 0.002 | 0.001 | 0.001 | **0.016** | **0.016** | **0.016** | **0.016** | **0.016** | **0.016** | **0.016** | **0.016** | **0.01** | **0.01** | ** | |
| Diptera sp. | OTU1255 | 0.016 | 0.021 | 0.015 | 0.019 | 0.021 | 0.019 | 0.017 | 0.018 | 0.014 | 0.017 | 0.14 | 0.14 | 0.14 | 0.14 | 0.14 | 0.14 | 0.14 | 0.14 | 0.14 | 0.14 | N.S. | |
| Calobatinae sp. | OTU1257 | 0.016 | 0.016 | 0.018 | 0.02 | 0.017 | 0.012 | 0.023 | 0.014 | 0.014 | 0.014 | 0.126 | 0.126 | 0.126 | 0.126 | 0.126 | 0.12 | 0.126 | 0.126 | 0.126 | 0.126 | N.S. | |
| Rhinophora lepida | OTU134 | 0.02 | 0.015 | 0.01 | 0.016 | 0.015 | 0.013 | 0.019 | 0.009 | 0.017 | 0.025 | 0.105 | 0.105 | 0.09 | 0.105 | 0.105 | 0.104 | 0.105 | 0.09 | 0.105 | 0.105 | N.S. | |
| Temnothorax affinis | OTU1437 | 0.015 | 0.016 | 0.017 | 0.023 | 0.025 | 0.012 | 0.014 | 0.011 | 0.018 | 0.015 | 0.112 | 0.112 | 0.112 | 0.112 | 0.112 | 0.11 | 0.112 | 0.11 | 0.112 | 0.112 | N.S. | |
| Diptera sp. | OTU1440 | 0.01 | 0.006 | 0.008 | 0.004 | 0.013 | 0.006 | 0.006 | 0.008 | 0.005 | 0.007 | **0.048** | **0.048** | **0.048** | **0.04** | **0.048** | **0.048** | **0.048** | **0.048** | **0.045** | **0.048** | * | |
| Aulonothroscus brevicollis | OTU1501 | 0.008 | 0.004 | 0.003 | 0.005 | 0.006 | 0.01 | 0.007 | 0.009 | 0.009 | 0.009 | **0.042** | **0.036** | **0.03** | **0.04** | **0.042** | **0.042** | **0.042** | **0.042** | **0.042** | **0.042** | * | |
| Sphegina clunipes | OTU1561 | 0.019 | 0.01 | 0.011 | 0.009 | 0.014 | 0.013 | 0.015 | 0.013 | 0.01 | 0.018 | 0.09 | 0.09 | 0.09 | 0.09 | 0.09 | 0.09 | 0.09 | 0.09 | 0.09 | 0.09 | N.S. | |
| Diptera sp. | OTU170 | 0.043 | 0.047 | 0.044 | 0.042 | 0.043 | 0.046 | 0.038 | 0.034 | 0.031 | 0.037 | 0.31 | 0.31 | 0.31 | 0.31 | 0.31 | 0.31 | 0.31 | 0.31 | 0.31 | 0.31 | N.S. | |
| Peripsocus cf. didymus | OTU1988 | 0.02 | 0.013 | 0.019 | 0.019 | 0.018 | 0.019 | 0.015 | 0.013 | 0.014 | 0.016 | 0.13 | 0.13 | 0.13 | 0.13 | 0.13 | 0.13 | 0.13 | 0.13 | 0.13 | 0.13 | N.S. | |
| Philotarsus picicornis | OTU2018 | 0.007 | 0.006 | 0.005 | 0.006 | 0.006 | 0.005 | 0.008 | 0.008 | 0.005 | 0.007 | **0.05** | **0.05** | **0.05** | **0.05** | **0.05** | **0.05** | **0.05** | **0.05** | **0.05** | **0.05** | * | |
| Chironomus luridus | OTU2093 | 0.02 | 0.016 | 0.018 | 0.018 | 0.015 | 0.014 | 0.02 | 0.019 | 0.016 | 0.021 | 0.14 | 0.14 | 0.14 | 0.14 | 0.14 | 0.14 | 0.14 | 0.14 | 0.14 | 0.14 | N.S. | |
| Diptera sp. | OTU2365 | 0.012 | 0.023 | 0.015 | 0.018 | 0.015 | 0.025 | 0.019 | 0.019 | 0.017 | 0.012 | 0.12 | 0.12 | 0.12 | 0.12 | 0.12 | 0.12 | 0.12 | 0.12 | 0.12 | 0.12 | N.S. | |
| Diptera sp. | OTU2616 | 0.013 | 0.017 | 0.019 | 0.012 | 0.018 | 0.026 | 0.016 | 0.021 | 0.016 | 0.018 | 0.12 | 0.128 | 0.128 | 0.12 | 0.128 | 0.128 | 0.128 | 0.128 | 0.128 | 0.128 | N.S. | |
| Megaselia verna | OTU2713 | 0.011 | 0.009 | 0.013 | 0.016 | 0.013 | 0.009 | 0.008 | 0.012 | 0.013 | 0.011 | 0.081 | 0.081 | 0.081 | 0.081 | 0.081 | 0.081 | 0.08 | 0.081 | 0.081 | 0.081 | N.S. | |
| Apodesmia incisula | OTU2966 | 0.013 | 0.01 | 0.014 | 0.014 | 0.013 | 0.016 | 0.011 | 0.01 | 0.012 | 0.014 | 0.1 | 0.1 | 0.1 | 0.1 | 0.1 | 0.1 | 0.1 | 0.1 | 0.1 | 0.1 | N.S. | |
| Hymenoptera sp. | OTU308 | 0.005 | 0.004 | 0.003 | 0.002 | 0.006 | 0.005 | 0.003 | 0.004 | 0.007 | 0.007 | **0.028** | **0.028** | **0.027** | **0.02** | **0.028** | **0.028** | **0.027** | **0.028** | **0.028** | **0.028** | * | |
| Herminia tarsipennalis | OTU333 | 0.012 | 0.023 | 0.017 | 0.019 | 0.015 | 0.026 | 0.017 | 0.016 | 0.013 | 0.019 | 0.12 | 0.12 | 0.12 | 0.12 | 0.12 | 0.12 | 0.12 | 0.12 | 0.12 | 0.12 | N.S. | |
| Coenosia sallae | OTU348 | 0.003 | 0.002 | 0.004 | 0.007 | 0.007 | 0.005 | 0.005 | 0.005 | 0.004 | 0.006 | **0.027** | **0.02** | **0.032** | **0.032** | **0.032** | **0.032** | **0.032** | **0.032** | **0.032** | **0.032** | * to ** | |
| Diptera sp. | OTU371 | 0.03 | 0.013 | 0.017 | 0.024 | 0.021 | 0.021 | 0.015 | 0.026 | 0.023 | 0.028 | 0.147 | 0.13 | 0.136 | 0.147 | 0.147 | 0.147 | 0.135 | 0.147 | 0.147 | 0.147 | N.S. | |
| Eilema sp. | OTU39 | 0.001 | 0.001 | 0.001 | 0.001 | 0.001 | 0.001 | 0.001 | 0.001 | 0.001 | 0.001 | **0.01** | **0.01** | **0.01** | **0.01** | **0.01** | **0.01** | **0.01** | **0.01** | **0.01** | **0.01** | ** | |
| Peromyia sp. | OTU472 | 0.003 | 0.005 | 0.004 | 0.004 | 0.008 | 0.006 | 0.005 | 0.004 | 0.005 | 0.005 | **0.03** | **0.036** | **0.036** | **0.036** | **0.036** | **0.036** | **0.036** | **0.036** | **0.036** | **0.036** | * | |
| Hymenoptera sp. | OTU474 | 0.003 | 0.003 | 0.003 | 0.003 | 0.002 | 0.002 | 0.003 | 0.002 | 0.003 | 0.002 | **0.02** | **0.02** | **0.02** | **0.02** | **0.02** | **0.02** | **0.02** | **0.02** | **0.02** | **0.02** | ** | |
| Platylabus sp. | OTU647 | 0.017 | 0.01 | 0.012 | 0.02 | 0.009 | 0.008 | 0.02 | 0.011 | 0.011 | 0.011 | 0.081 | 0.081 | 0.081 | 0.081 | 0.081 | 0.08 | 0.081 | 0.081 | 0.081 | 0.081 | N.S. | |
| Stenomacrus celer | OTU710 | 0.005 | 0.005 | 0.004 | 0.009 | 0.005 | 0.004 | 0.003 | 0.003 | 0.006 | 0.004 | **0.032** | **0.032** | **0.032** | **0.032** | **0.032** | **0.032** | **0.03** | **0.03** | **0.032** | **0.032** | * | |
| Phaonia sp. | OTU754 | 0.018 | 0.009 | 0.022 | 0.023 | 0.032 | 0.019 | 0.028 | 0.025 | 0.025 | 0.031 | 0.162 | 0.09 | 0.162 | 0.162 | 0.162 | 0.162 | 0.162 | 0.162 | 0.162 | 0.162 | N.S. | |
| Diptera sp. | OTU80 | 0.007 | 0.004 | 0.011 | 0.007 | 0.013 | 0.006 | 0.008 | 0.008 | 0.009 | 0.012 | 0.056 | **0.04** | 0.056 | 0.056 | 0.056 | 0.054 | 0.056 | 0.056 | 0.056 | 0.056 | N.S. to * | |
| Cicadellidae sp. | OTU868 | 0.001 | 0.001 | 0.001 | 0.001 | 0.001 | 0.001 | 0.001 | 0.001 | 0.001 | 0.001 | 0.01 | 0.01 | 0.01 | 0.01 | 0.01 | 0.01 | 0.01 | 0.01 | 0.01 | 0.01 | ** | |
| Lasius platythorax | OTU870 | 0.014 | 0.015 | 0.021 | 0.029 | 0.019 | 0.011 | 0.016 | 0.014 | 0.024 | 0.017 | 0.126 | 0.126 | 0.126 | 0.126 | 0.126 | 0.11 | 0.126 | 0.126 | 0.126 | 0.126 | N.S. | |
| Glyphipterix forsterella | OTU922 | 0.005 | 0.017 | 0.02 | 0.019 | 0.019 | 0.011 | 0.01 | 0.02 | 0.016 | 0.012 | **0.05** | 0.096 | 0.096 | 0.096 | 0.096 | 0.09 | 0.09 | 0.096 | 0.096 | 0.09 | N.S. to * | |
| Lethades facialis | OTU1047 | 0.015 | 0.016 | 0.015 | 0.017 | 0.014 | 0.02 | 0.019 | 0.013 | 0.015 | 0.017 | 0.13 | 0.13 | 0.13 | 0.13 | 0.13 | 0.13 | 0.13 | 0.13 | 0.13 | 0.13 | N.S. | MEDIUM |
| Compsilura concinnata | OTU1108 | 0.011 | 0.007 | 0.006 | 0.003 | 0.002 | 0.008 | 0.003 | 0.006 | 0.004 | 0.005 | **0.03** | **0.03** | **0.03** | **0.027** | **0.02** | **0.03** | **0.027** | **0.03** | **0.028** | **0.03** | * | |
| Argyresthia conjugella | OTU1193 | 0.01 | 0.017 | 0.019 | 0.025 | 0.011 | 0.013 | 0.013 | 0.011 | 0.015 | 0.008 | 0.09 | 0.09 | 0.09 | 0.09 | 0.09 | 0.09 | 0.09 | 0.09 | 0.09 | 0.08 | N.S. | |
| Diptera sp. | OTU1300 | 0.033 | 0.035 | 0.037 | 0.025 | 0.034 | 0.028 | 0.037 | 0.037 | 0.046 | 0.029 | 0.252 | 0.252 | 0.252 | 0.25 | 0.252 | 0.252 | 0.252 | 0.252 | 0.252 | 0.252 | N.S. | |
| Lonchaea postica | OTU1320 | 0.02 | 0.031 | 0.027 | 0.024 | 0.018 | 0.028 | 0.023 | 0.025 | 0.032 | 0.027 | 0.18 | 0.184 | 0.184 | 0.184 | 0.18 | 0.184 | 0.184 | 0.184 | 0.184 | 0.184 | N.S. | |
| Stromboceros delicatulus | OTU1366 | 0.015 | 0.015 | 0.012 | 0.01 | 0.012 | 0.009 | 0.016 | 0.01 | 0.008 | 0.007 | 0.072 | 0.072 | 0.072 | 0.072 | 0.072 | 0.072 | 0.072 | 0.072 | 0.072 | 0.07 | N.S. | |
| Passaloecus insignis | OTU1504 | 0.044 | 0.041 | 0.035 | 0.043 | 0.046 | 0.039 | 0.045 | 0.039 | 0.034 | 0.038 | 0.34 | 0.34 | 0.34 | 0.34 | 0.34 | 0.34 | 0.34 | 0.34 | 0.34 | 0.34 | N.S. | |
| Tipula scripta | OTU1535 | 0.014 | 0.013 | 0.017 | 0.016 | 0.015 | 0.022 | 0.027 | 0.015 | 0.01 | 0.011 | 0.104 | 0.104 | 0.104 | 0.104 | 0.104 | 0.104 | 0.104 | 0.104 | 0.1 | 0.1 | N.S. | |
| Perilissus rufoniger | OTU1671 | 0.021 | 0.012 | 0.018 | 0.015 | 0.02 | 0.022 | 0.022 | 0.017 | 0.021 | 0.017 | 0.136 | 0.12 | 0.136 | 0.135 | 0.136 | 0.136 | 0.136 | 0.136 | 0.136 | 0.136 | N.S. | |
| Hylaeus confusus | OTU2070 | 0.019 | 0.014 | 0.021 | 0.021 | 0.019 | 0.017 | 0.018 | 0.02 | 0.019 | 0.029 | 0.153 | 0.14 | 0.153 | 0.153 | 0.153 | 0.153 | 0.153 | 0.153 | 0.153 | 0.153 | N.S. | |
| Diplazon sp. | OTU2274 | 0.012 | 0.027 | 0.016 | 0.015 | 0.012 | 0.023 | 0.017 | 0.018 | 0.019 | 0.019 | 0.12 | 0.12 | 0.12 | 0.12 | 0.12 | 0.12 | 0.12 | 0.12 | 0.12 | 0.12 | N.S. | |
| Trichosia edwardsi | OTU2607 | 0.007 | 0.006 | 0.006 | 0.002 | 0.007 | 0.008 | 0.002 | 0.006 | 0.001 | 0.008 | **0.042** | **0.042** | **0.042** | **0.018** | **0.042** | **0.042** | **0.018** | **0.042** | **0.01** | **0.042** | * to ** | |
| Lyciella rorida | OTU261 | 0.038 | 0.024 | 0.029 | 0.026 | 0.039 | 0.029 | 0.027 | 0.027 | 0.03 | 0.034 | 0.24 | 0.24 | 0.24 | 0.24 | 0.24 | 0.24 | 0.24 | 0.24 | 0.24 | 0.24 | N.S. | |
| Eustalomyia festiva | OTU278 | 0.025 | 0.016 | 0.01 | 0.021 | 0.015 | 0.02 | 0.016 | 0.023 | 0.021 | 0.025 | 0.135 | 0.135 | 0.1 | 0.135 | 0.135 | 0.135 | 0.135 | 0.135 | 0.135 | 0.135 | N.S. | |
| Scathophaga sp. | OTU292 | 0.043 | 0.029 | 0.034 | 0.039 | 0.027 | 0.026 | 0.024 | 0.032 | 0.033 | 0.027 | 0.24 | 0.24 | 0.24 | 0.24 | 0.24 | 0.24 | 0.24 | 0.24 | 0.24 | 0.24 | N.S. | |
| Macquartia dispar | OTU318 | 0.017 | 0.016 | 0.022 | 0.024 | 0.025 | 0.024 | 0.028 | 0.019 | 0.016 | 0.022 | 0.16 | 0.16 | 0.16 | 0.16 | 0.16 | 0.16 | 0.16 | 0.16 | 0.16 | 0.16 | N.S. | |
| Leptosciarella scutellata | OTU322 | 0.019 | 0.026 | 0.021 | 0.024 | 0.027 | 0.018 | 0.017 | 0.025 | 0.022 | 0.017 | 0.17 | 0.17 | 0.17 | 0.17 | 0.17 | 0.17 | 0.17 | 0.17 | 0.17 | 0.17 | N.S. | |
| Melinda sp. | OTU349 | 0.021 | 0.037 | 0.026 | 0.038 | 0.047 | 0.037 | 0.036 | 0.027 | 0.033 | 0.042 | 0.21 | 0.234 | 0.234 | 0.234 | 0.234 | 0.234 | 0.234 | 0.234 | 0.234 | 0.234 | N.S. | |
| Volucella sp. | OTU420 | 0.005 | 0.007 | 0.006 | 0.005 | 0.002 | 0.007 | 0.008 | 0.003 | 0.006 | 0.006 | **0.04** | **0.04** | **0.04** | **0.04** | **0.02** | **0.04** | **0.04** | **0.027** | **0.04** | **0.04** | * | |
| Syrphinae sp. | OTU44 | 0.014 | 0.016 | 0.016 | 0.013 | 0.016 | 0.009 | 0.017 | 0.016 | 0.019 | 0.018 | 0.117 | 0.117 | 0.117 | 0.117 | 0.117 | 0.09 | 0.117 | 0.117 | 0.117 | 0.117 | N.S. | |
| Tenthredo sp. | OTU442 | 0.013 | 0.01 | 0.009 | 0.015 | 0.008 | 0.008 | 0.012 | 0.012 | 0.01 | 0.007 | 0.072 | 0.072 | 0.072 | 0.072 | 0.072 | 0.072 | 0.072 | 0.072 | 0.072 | 0.07 | N.S. | |
| Phaonia angelicae | OTU47 | 0.002 | 0.003 | 0.002 | 0.002 | 0.002 | 0.001 | 0.003 | 0.001 | 0.001 | 0.003 | 0.014 | 0.014 | 0.014 | 0.014 | 0.014 | 0.01 | 0.014 | 0.01 | 0.01 | 0.014 | ** | |
| Phaonia rufiventris | OTU495 | 0.019 | 0.017 | 0.022 | 0.018 | 0.014 | 0.018 | 0.013 | 0.014 | 0.013 | 0.01 | 0.117 | 0.117 | 0.117 | 0.117 | 0.117 | 0.117 | 0.117 | 0.117 | 0.117 | 0.1 | N.S. | |
| Hymenoptera sp. | OTU660 | 0.004 | 0.004 | 0.004 | 0.002 | 0.006 | 0.005 | 0.006 | 0.004 | 0.004 | 0.009 | **0.036** | **0.036** | **0.036** | **0.02** | **0.036** | **0.036** | **0.036** | **0.036** | **0.036** | **0.036** | * to ** | |
| Pieris sp. | OTU688 | 0.012 | 0.02 | 0.014 | 0.014 | 0.019 | 0.016 | 0.018 | 0.015 | 0.013 | 0.024 | 0.12 | 0.12 | 0.12 | 0.12 | 0.12 | 0.12 | 0.12 | 0.12 | 0.12 | 0.12 | N.S. | |
| Eristalis pertinax | OTU725 | 0.015 | 0.016 | 0.023 | 0.023 | 0.028 | 0.014 | 0.028 | 0.021 | 0.016 | 0.022 | 0.14 | 0.14 | 0.14 | 0.14 | 0.14 | 0.14 | 0.14 | 0.14 | 0.14 | 0.14 | N.S. | |
| Suillia ustulata | OTU729 | 0.027 | 0.031 | 0.03 | 0.035 | 0.029 | 0.038 | 0.03 | 0.021 | 0.03 | 0.034 | 0.243 | 0.243 | 0.243 | 0.243 | 0.243 | 0.243 | 0.243 | 0.21 | 0.243 | 0.243 | N.S. | |
| Bradysia placida | OTU826 | 0.029 | 0.035 | 0.044 | 0.048 | 0.047 | 0.045 | 0.043 | 0.049 | 0.049 | 0.047 | 0.29 | 0.315 | 0.344 | 0.344 | 0.344 | 0.344 | 0.344 | 0.344 | 0.344 | 0.344 | N.S. | |
| Muscina pascuorum | OTU937 | 0.014 | 0.017 | 0.013 | 0.017 | 0.013 | 0.017 | 0.02 | 0.016 | 0.02 | 0.013 | 0.13 | 0.13 | 0.13 | 0.13 | 0.13 | 0.13 | 0.13 | 0.13 | 0.13 | 0.13 | N.S. | |
| Dusona sp. | OTU997 | 0.023 | 0.024 | 0.016 | 0.019 | 0.013 | 0.022 | 0.019 | 0.011 | 0.016 | 0.025 | 0.128 | 0.128 | 0.128 | 0.128 | 0.117 | 0.128 | 0.128 | 0.11 | 0.128 | 0.128 | N.S. | |
| Polistes sp. | OTU1093 | 0.013 | 0.011 | 0.014 | 0.013 | 0.011 | 0.013 | 0.013 | 0.011 | 0.007 | 0.016 | 0.099 | 0.099 | 0.099 | 0.099 | 0.099 | 0.099 | 0.099 | 0.099 | 0.07 | 0.099 | N.S. | HIGH |
| Dichrogaster sp. | OTU1104 | 0.032 | 0.03 | 0.033 | 0.025 | 0.033 | 0.034 | 0.033 | 0.036 | 0.036 | 0.038 | 0.27 | 0.27 | 0.27 | 0.25 | 0.27 | 0.27 | 0.27 | 0.27 | 0.27 | 0.27 | N.S. | |
| Raphidioptera | OTU1229 | 0.038 | 0.033 | 0.038 | 0.038 | 0.041 | 0.05 | 0.037 | 0.02 | 0.034 | 0.035 | 0.297 | 0.297 | 0.297 | 0.297 | 0.297 | 0.297 | 0.297 | 0.2 | 0.297 | 0.297 | N.S. | |
| Diptera sp. | OTU1471 | 0.036 | 0.032 | 0.037 | 0.036 | 0.036 | 0.024 | 0.033 | 0.036 | 0.032 | 0.03 | 0.27 | 0.27 | 0.27 | 0.27 | 0.27 | 0.24 | 0.27 | 0.27 | 0.27 | 0.27 | N.S. | |
| Spelobia quinata | OTU1954 | 0.006 | 0.01 | 0.014 | 0.01 | 0.018 | 0.013 | 0.012 | 0.006 | 0.011 | 0.012 | 0.06 | 0.08 | 0.08 | 0.08 | 0.08 | 0.08 | 0.08 | 0.06 | 0.08 | 0.08 | N.S. | |
| Lymnophyes sp. | OTU2297 | 0.038 | 0.026 | 0.05 | 0.033 | 0.041 | 0.041 | 0.027 | 0.04 | 0.032 | 0.032 | 0.26 | 0.26 | 0.26 | 0.26 | 0.26 | 0.26 | 0.26 | 0.26 | 0.26 | 0.26 | N.S. | |
| Scaeva sp. | OTU26 | 0.038 | 0.039 | 0.044 | 0.039 | 0.041 | 0.037 | 0.037 | 0.035 | 0.034 | 0.037 | 0.34 | 0.34 | 0.34 | 0.34 | 0.34 | 0.34 | 0.34 | 0.34 | 0.34 | 0.34 | N.S. | |
| Aphantopus hyperantus | OTU876 | 0.039 | 0.039 | 0.039 | 0.038 | 0.039 | 0.037 | 0.044 | 0.037 | 0.039 | 0.035 | 0.35 | 0.35 | 0.35 | 0.35 | 0.35 | 0.35 | 0.35 | 0.35 | 0.35 | 0.35 | N.S. | |
